# *DNMT3A* knockouts in human iPSCs prevent *de novo* DNA methylation and reveal growth advantage during hematopoietic differentiation

**DOI:** 10.1101/2021.12.15.472782

**Authors:** Olivia Cypris, Julia Franzen, Joana Frobel, Philipp Glück, Chao-Chung Kuo, Stephani Schmitz, Selina Nüchtern, Martin Zenke, Wolfgang Wagner

## Abstract

DNA methyltransferase 3A (*DNMT3A*) is a frequently mutated gene in many hematological malignancies, indicating that it may be essential for hematopoietic differentiation. Here, we addressed the functional relevance of DNMT3A for differentiation of human induced pluripotent stem cells (iPSCs) by knocking out exon 2, 19, or 23. Exon 19^-/-^ and 23^-/-^ lines revealed absence of almost the entire *de novo* DNA methylation during mesenchymal and hematopoietic differentiation. Yet, differentiation was only slightly reduced in exon 19^-/-^ and increased in exon 23^-/-^ lines, whereas there was no significant impact on gene expression in hematopoietic progenitors (iHPCs). Notably, *DNMT3A*^-/-^ iHPCs recapitulate some DNA methylation differences of acute myeloid leukemia with *DNMT3A* mutations. Furthermore, multicolor genetic barcoding revealed competitive growth advantage of exon 23^-/-^ iHPCs. Our results demonstrate that *de novo* DNA methylation during hematopoietic differentiation of iPSCs is almost entirely dependent on DNMT3A and exon 23^-/-^ iHPCs even gained growth advantage.

## Introduction

DNA methylation (DNAm) is tightly regulated during cellular differentiation (Meissner et al., 2008). *De novo* methylation is primarily established by two DNA methyltransferases: DNMT3A and DNMT3B (Okano et al., 1999). The latter is rather active in early development and specifically methylates minor satellite repeats, while DNMT3A was suggested to be more important in later development (Okano *et al*., 1999). DNMT3A has five different protein-coding splice variants which might have different functional roles and are spliced in tissue and disease specific manner (Bozic et al., 2018; Lin et al., 2017). So far, it is largely unclear how DNAm is targeted to specific sites in the genome.

DNMT3A seems to be of particular relevance for hematopoietic differentiation, since it frequently reveals heterozygous mutations in acute myeloid leukemia (AML) and other hematological malignancies (Ley et al., 2010; Roller et al., 2013). In AML, about 65% of the *DNMT3A* mutations are located in the hotspot R882 in exon 23 (Brunetti et al., 2017; Emperle et al., 2019). This mutation is believed to reduce enzyme activity by blocking its ability to form active tetramers with the wildtype form (Russler-Germain et al., 2014) and to induce aberrant DNAm patterns due to conformational changes in the enzyme (Emperle *et al*., 2019). Furthermore, *DNMT3A* holds the most frequent driver mutations in clonal hematopoiesis of indeterminate potential (CHIP) (Challen and Goodell, 2020).

The functional relevance of Dnmt3a was investigated in homozygous knockout mice already more than 20 years ago, demonstrating that development remained possible despite undersize at birth and death at about 4 weeks of age (Okano *et al*., 1999). More recent analysis of murine *Dnmt3a* knockout models revealed an increased self-renewal capacity of hematopoietic stem cells (HSCs) with reduced differentiation capacity (Jeong et al., 2018). Heterozygous knockouts had only a subtle effect on methylation in mice with a bias for myeloid differentiation and higher propensity for malignant transformation (Cole et al., 2017). Other studies reported that knocking out *Dnmt3a* in HSCs and transplanting them into mice leads to development of a wide range of myeloid and lymphoid malignancies (Celik et al., 2015; Mayle et al., 2015; Peters et al., 2014).

In contrast, relatively few studies addressed the consequences of *DNMT3A* knockout in human cells. In human embryonic stem cells no immediate negative effects of DNMT3A^-/-^ could be observed on downstream differentiation, although global loss of DNA methylation was observed after many passages with focal areas of hypermethylation (Liao et al., 2015). Recently, it has been demonstrated that human induced pluripotent stem cells (iPSCs) with knockout in *DNMT3A* can be differentiated into cardiomyocytes with similar efficiency as wildtype controls (Madsen et al., 2020). However, these iPSC-derived cardiomyocytes showed global hypomethylation with differences in contractile behavior and an aberrant activation of glucose and lipid metabolism (Madsen *et al*., 2020).

In this study, we investigated the functional relevance of DNMT3A for hematopoietic differentiation of human iPSCs. To this end, we generated iPSC lines with knockout of exon 2, which contains a start codon for transcripts 1, 3 and 4. Alternatively, we removed exon 19, which has also been targeted in the above-mentioned studies, or exon 23 that comprises the frequently mutated hotspot R882. We then analyzed the effect on growth and differentiation of iPSCs toward mesenchymal and hematopoietic lineages as well as its impact on DNA methylation and gene expression and whether clones gained growth advantage over wildtype cells.

## Results

### Knockout of *DNMT3A* exons does not evoke phenotypic changes in human iPSCs

To generate knockouts of different *DNMT3A* exons, we used iPSC lines from three donors for CRIPSR/Cas9n targeting of the intron/exon boundaries of exon 2 and exon 19, or excision of exon 23 (Fig. 1a; Supplemental file 1: Fig. S1). For exon 2, we achieved a homozygous truncation of *DNMT3A* for donor 1, due to a new in frame start codon in exon 3, and a heterozygous knockout for donor 2. Furthermore, we generated two homozygous knockout lines for exon 19 and two homozygous knockout lines for exon 23, as determined by Western blot analysis (Fig. 1b). These results were further substantiated by quantitative RT-PCR, indicating that knockout of exon 23 still enabled truncated transcripts (Fig. 1c).

**Fig. 1:**
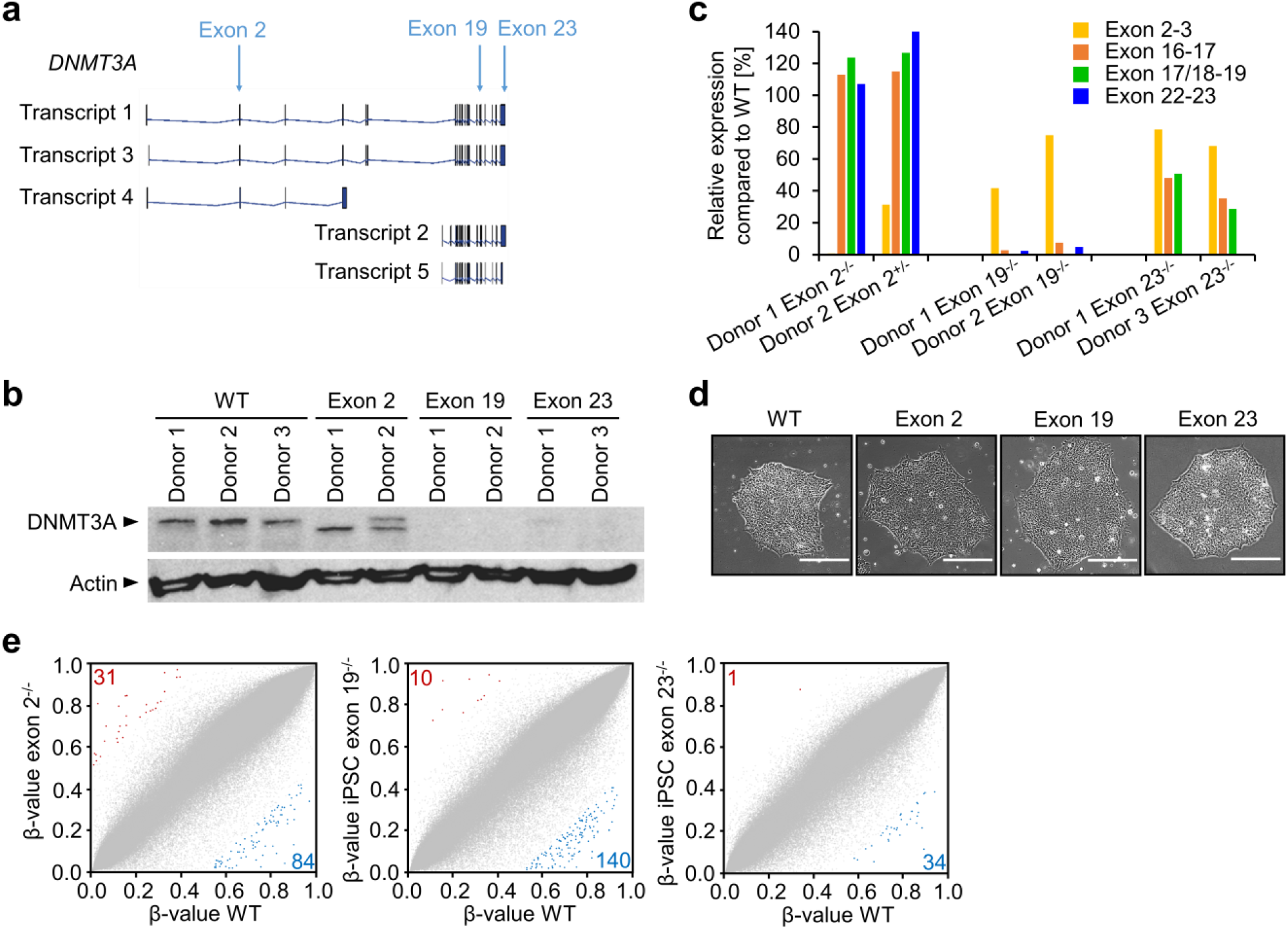
Characterization of different *DNMT3A* exon knockouts in iPSCs. **a)** Schematic overview of the protein coding *DNMT3A* splice variants. Arrows indicate CRISPR/Cas9n target sites. **b)** Western blot analysis of DNMT3A upon knockout of exon 2, exon 19, or exon 23. Arrow corresponds to height of transcript 1 / 3. WT = wildtype. **c)** Semiquantitative RT-PCR analysis of expression of *DNMT3A* exons (normalized to *GAPDH*, measured in duplicate). **d)** Representative phase contrast images of iPSC colonies with *DNMT3A* knockouts and WT. Scale bar = 150 µm. **e)** Scatter plots show DNAm levels (β-values) for all measured CpG sites (grey) when comparing WT *versus* knockout of exon 2, 19, or 23, respectively. For exon 2, we only present results for the homozygous knockout in exon 2, whereas for exon 19 and 23 the mean of two homozygous knockout lines is presented. CpGs with more than 50% hypermethylation (red) or hypomethylation (blue) are indicated.

*DNMT3A* knockouts did not reveal negative effects on growth and differentiation of iPSCs. All clones showed typical iPSC-like morphology (Fig. 1d), expressed the pluripotency markers OCT3/4 and TRA1-60, and were able to differentiate toward endodermal, mesodermal and ectodermal lineages (Supplemental file 1: Fig. S2a,b). To assess the impact of DNMT3A loss on DNA methylation, we used the Infinium Methylation EPIC BeadChip technology. Multidimensional scaling (MDS) indicated that DNAm profiles were similar in wildtype and *DNMT3A* knockout iPSC lines (Supplemental file 1: Fig. S2c). Pairwise comparison showed that only few CpGs had a clear conversion of DNAm levels (at least 50% gain or loss in DNAm) upon knockout, while there was no systematic off-set in DNAm levels (Fig. 1e, Supplemental file 2: Tab. S5). Taken together, we were able to successfully knock out three different exons of *DNMT3A* in human iPSCs and this did not evoke pronounced effects on DNAm levels while in a pluripotent cell state.

### *DNMT3A* knockouts impair *de novo* DNAm in iPSC-derived mesenchymal cells

To test the impact of *DNMT3A* knockouts on directed differentiation, we initially differentiated all iPSC lines toward mesenchymal stromal cells (iMSCs) that constitute an important part of the hematopoietic niche. After five weeks, differentiated cells revealed typical fibroblastoid morphology (Fig. 2a) and upregulation of immunophenotypic markers for MSCs, albeit expression of CD105 and CD73 was lower in exon 19^-/-^ clones (Fig. 2b, Supplemental file 1: Fig. S3a). The differentiation of wildtype iPSCs towards iMSCs was associated with pronounced DNAm changes of more than 50% at many CpGs: wildtype iMSCs gained DNAm at 2 135 CpGs and lost methylation at 6 700 CpGs (Fig. 2c; Supplemental file 3: Tab. S6). Notably, hypermethylated CpGs were enriched in promoter regions of genes in functional categories associated with mesenchymal differentiation, such as skeletal system development, osteoblast development, and cartilage development, indicating that *de novo* DNAm during iMSC differentiation is indeed lineage specific (Supplemental file 1: Fig. S3b). A similar number of differentially methylated CpGs were observed for the exon 2^-/-^ clone (2 509 hyper- and 9 771 hypomethylated CpGs) and most of them were overlapping with the DNAm changes of wildtype clones (Supplemental file 1: Fig. S3c). This might be anticipated, due to a new start codon in exon 3 maintaining the functional MTase-domain. In contrast, exon 19^-/-^ and exon 23^-/-^ clones hardly gained DNAm during differentiation towards iMSCs (only 31 and 32 CpGs, respectively), while hypomethylation followed a similar pattern as in wildtype iPSCs (10 424 and 5 857 CpGs, respectively). In fact, there was a very high overlap of hypomethylated CpGs in wildtype, exon 2^-/-^, exon 19 ^-/-^, and exon 23^-/-^ cells (Supplemental file 1: Fig. S3c). Our results demonstrate that despite phenotypic similarities of iMSCs there was hardly any *de novo* DNAm during differentiation of iPSCs with knockout of *DNMT3A* exon 19 and exon 23 toward the mesenchymal lineage.

**Fig. 2:**
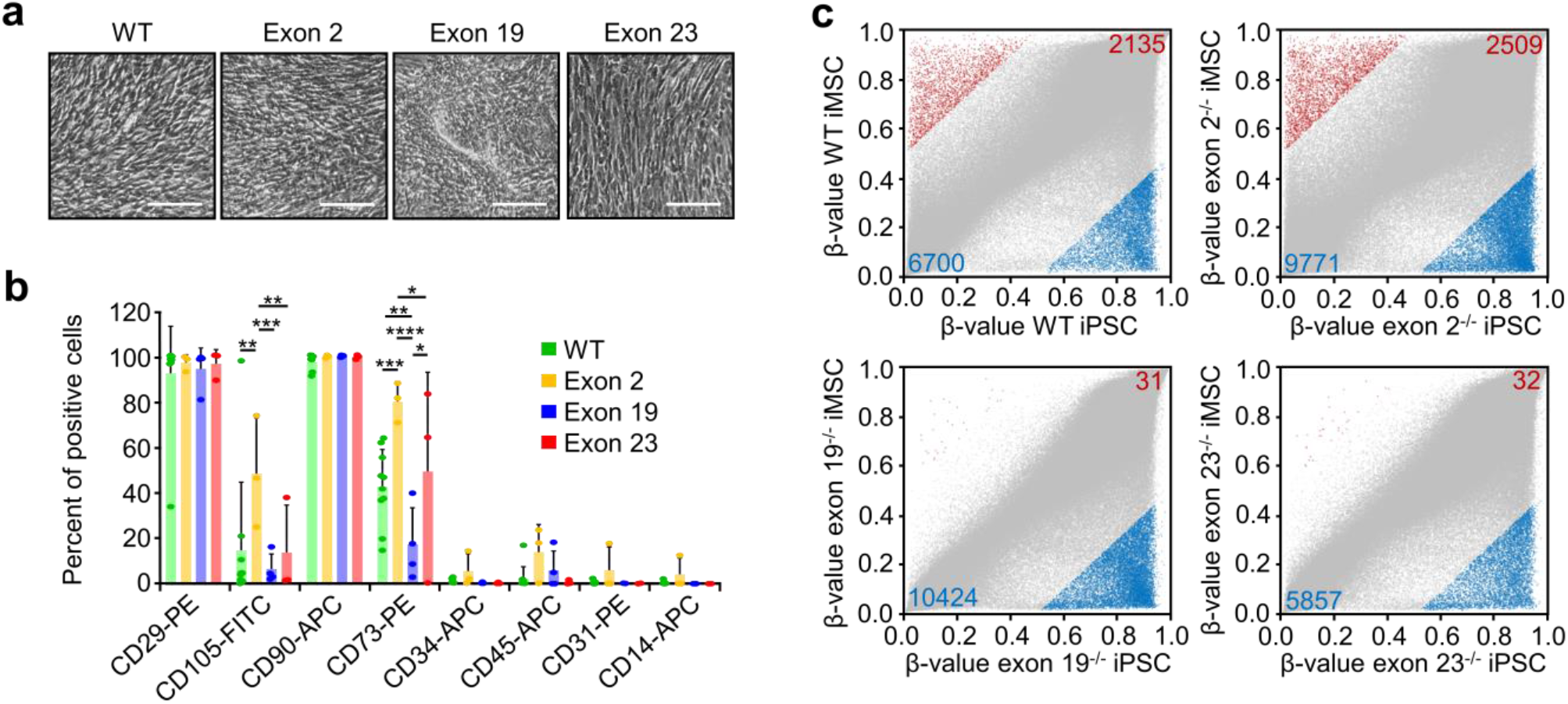
Characterization of iPSC-derived mesenchymal stromal cells. **a)** Representative phase contrast images of iPSC-derived mesenchymal stromal cells (iMSCs) at day 35 for wildtype (WT) and knockout lines. Scale bar = 250 µm. **b)** Flow cytometric analysis of surface markers of iMSCs at day 35. Significance was estimated by 2-way ANOVA with Tukey’s PostHoc Test; n = 10 for WT, n = 3 for exon 2 (exon 2^-/-^ and exon 2^+/-^), n = 4 for exon 19, and n = 3 for exon 23 (* *P* < 0.05; ** *P* < 0.01, *** *P* < 0.001, **** *P* < 0.0001; mean ± SD). **c)** Scatter plots of DNAm levels (β-values) for all measured CpG sites (grey) in iMSCs derived from homozygous knockout lines for exon 2 (n = 1), exon 19 (n = 2), or exon 23 (n = 2), in comparison to corresponding WT lines. CpGs with more than 50% hyper- (red) and hypomethylation (blue) are indicated. Darker blue and red spots represent promoter-associated CpGs.

### Hematopoietic differentiation is slightly decreased in exon 19^-/-^ and in tendency increased in exon 23^-/-^

Mutations in *DNMT3A* are frequently observed in hematopoietic malignancies and therefore we investigated if hematopoietic differentiation was affected in iPSCs with *DNMT3A* knockouts in exon 19 or 23 (Fig. 3a). We did not further consider the exon 2^-/-^ clone, since we did not have a homozygous biological replicate and no effects on DNAm were observed in iMSCs. After 16 days, all clones produced non-adherent cells with typical hematological morphology (Fig. 3b) with a higher frequency in exon 23^-/-^ clones (Fig. 3c). Furthermore, hematopoietic surface markers were upregulated for all clones, but CD31, CD33, CD34 and CD43 were less expressed in exon 19^-/-^ lines (Fig. 3d; Supplemental file 1: Fig. S4a), whereas CD61 and CD235α were particularly up-regulated in exon 23^-/-^ lines. We then tested the colony forming unit (CFU) potential in methylcellulose. In exon 19^-/-^ clones the CFU frequency was reduced and biased for CFU macrophage colonies (Fig. 3e; Supplemental file 1: Fig. S4b). Overall, all iPSC lines were capable of differentiating towards hematopoietic lineage, but propensity was reduced in exon 19^-/-^ as compared to wildtype or exon 23^-/-^ clones.

**Fig. 3:**
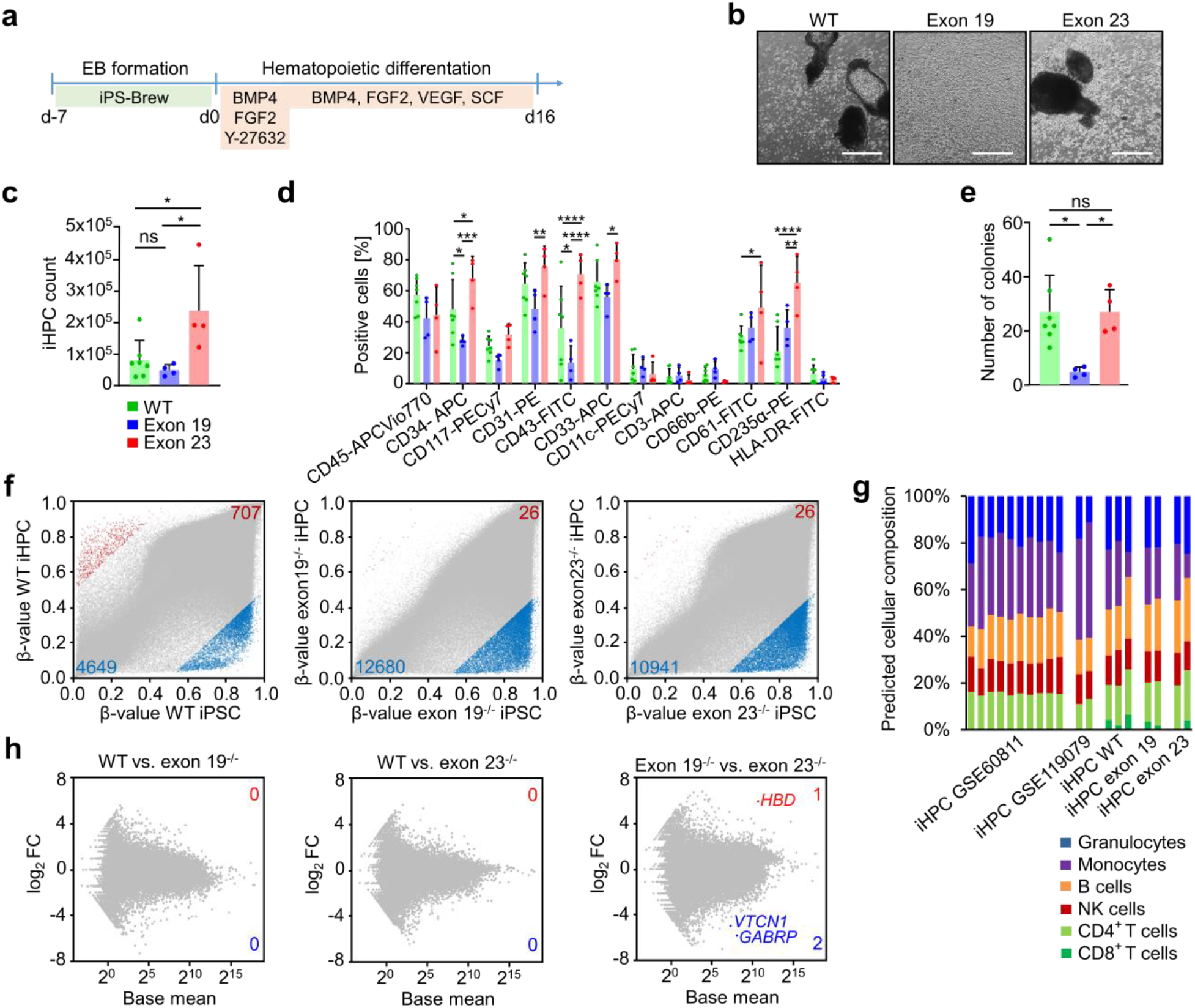
iPSC-derived hematopoietic progenitor cells. Schematic representation of the protocol used for hematopoietic differentiation of iPSCs. Representative phase contrast images of produced hematopoietic progenitor cells on d16 of hematopoietic differentiation. Scale bar = 500 µm. **c)** Absolute count of harvested iHPCs per well. Statistics were calculated with 1-way ANOVA and Tukey’s PostHoc test; n = 7 for wildtype (WT), n = 4 for each knockout (* *P* < 0.05; mean ± SD). **d)** Surface marker expression of hematopoietic progenitors. Statistics were calculated with 2-way ANOVA and Tukey’s PostHoc test; n = 7 for WT, n = 4 for each knockout (* *P* < 0.05; ** *P* < 0.01, *** *P* < 0.001, **** *P* < 0.0001; mean ± SD). **e)** Total number of colonies formed in CFU assays. Statistics were calculated with 1-way ANOVA and Tukey’s PostHoc test; n = 7 for WT, n = 4 for each knockout (* *P* < 0.05; mean ± SD). **f)** Scatter plots show β-values for all measured CpG sites (grey) when comparing the mean of wildtype iHPCs, exon 19^-/-^ iHPCs or exon 23^-/-^ iHPCs with their respective iPSC counterparts. Hypermethylated CpGs are depicted in red, hypomethylated CpGs in blue with delta mean β-value >0.5 or <-0.5. CpG sites associated with promotor regions are shown without transparency. **(g)** A deconvolution algorithm (Houseman *et al*., 2012) was used to estimate the composition of different mature hematopoietic cell types based on the DNAm profiles of our iHPCs, those of Nishizawa et al. (Nishizawa *et al*., 2016) and of our previously published iHPC data (Cypris *et al*., 2019). **h)** Gene expression changes compared between WT iHPCs and exon 19^-/-^ or exon 23^-/-^ iHPCs depicted as log_2_ fold change against the mean of normalized counts of all samples regarding sequencing depth (=base mean). Significantly differentially expressed genes are highlighted.

### Impaired *de novo* DNAm during hematopoietic differentiation is not reflected by the transcriptome

We subsequently analyzed DNAm changes after 16 days of hematopoietic differentiation. After differentiation, wildtype clones revealed strong DNAm changes (more than 50%) at many CpGs: 707 CpGs were hypermethylated and 4 649 CpGs hypomethylated. In contrast, only 26 CpGs became hypermethylated in *DNMT3A* exon 19^-/-^, while 12 680 CpGs became hypomethylated with the same thresholds. A similar effect was observed for *DNMT3A* exon 23^-/-^ lines (26 CpGs hyper- and 10 941 CpGs hypomethylated; Fig. 3f; Supplemental file 4: Tab. S7). The overlap of CpGs with pronounced gains and losses in DNAm was very high between wildtype and knockout lines (Supplemental file 1: Fig. S5a). Furthermore, wildtype *versus* knockout iHPC clones revealed much higher DNAm at many CpGs in wildtype, whereas this was hardly observed in exon 19^-/-^ or exon 23^-/-^ (Supplemental file 1: Fig. S5b; Supplemental file 5: Tab. S8). Thus, *DNMT3A* knockout in either exon 19 or exon 23 clearly impairs *de novo* DNAm during hematopoietic differentiation of iPSCs.

The moderate differences in hematopoietic differentiation between exon 19^-/-^ and exon 23^-/-^ clones might be attributed to different DNA methylation profiles. Overall, only 110 CpGs were more than 50% hypomethylated and 353 CpGs were more than 50% hypermethylated in exon 23^-/-^ *versus* exon 19^-/-^ lines. Very similar results were observed when we only compared the two samples of donor 1, to prevent impact of different genetic background (Supplemental file 1: Fig. S6a,b).

To estimate if *DNMT3A* knockouts affected differentiation toward specific hematopoietic lineages, we used a deconvolution algorithm to determine the composition of hematopoietic subsets based on DNAm profiles (Houseman et al., 2012) (Fig. 3g). DNAm profiles of wildtype iHPCs indicated multilineage differentiation, similar to previously published iHPC profiles (Cypris et al., 2019; Nishizawa et al., 2016). Despite the impact of exon 19^-/-^ and exon 23^-/-^ on *de novo* DNAm, estimates for cellular composition were similar between wildtype and knockout iHPCs. This might be attributed to the finding that cell-type specific DNAm patterns of such signatures are rather hypomethylated (Houseman *et al*., 2012; Schmidt et al., 2020) and such hypomethylation was largely maintained in *DNMT3A* knockouts. Alternatively, we focused on individual CpGs that are specifically hypomethylated in granulocytes, monocytes, B cells, CD4, CD8, or NK cells (Cypris *et* al., 2019; Frobel et al., 2018). Overall, DNAm patterns at these lineage-specific sites were similar in wildtype and *DNMT3A* knockout iHPCs (Supplemental file 1: Fig. S5c).

To investigate if impaired *de novo* DNAm in *DNMT3A* knockouts was also reflected on transcriptomic level, we performed RNA-sequencing analysis. Unexpectedly, there were no significant differences in wildtype *versus* either exon 19^-/-^ or exon 23^-/-^ clones (Wald test; *P* < 0.05; Fig. 3h; Supplemental file 6: Tab. S9). Furthermore, even those genes that showed a difference of more than 50% on DNAm levels did not reveal a clear tendency for differential expression, even if we focused on DNAm changes in the promoter regions (Supplemental file 1: Fig. S5d). When we compared differential gene expression of iHPCs between exon 19^-/-^ or exon 23^-/-^ clones, only three genes reached significance (Hemoglobin Subunit Delta (*HBD*), V-Set Domain Containing T Cell Activation Inhibitor 1 (*VTCN1*), and Gamma-Aminobutyric Acid Type A Receptor Subunit Pi (*GABRP*; Fig. 3h). Taken together, loss of *de novo* methylation in exon 19^-/-^ and exon 23^-/-^ clones was not reflected on differential gene expression.

### Impact of *DNMT3A* knockouts on specific DNAm changes

To gain better insight into how global DNAm patterns change in iMSCs and iHPCs, we performed a principal component analysis (PCA), demonstrating that despite the marked block of *de novo* DNAm, there are still cell-type specific changes, which can be particularly attributed to focal hypomethylation (Fig. 4a). Notably, many pluripotency-associated DNAm patterns, which are usually lost during differentiation, were maintained in iMSCs and iHPCs of exon 19^-/-^ and exon 23^-/-^ lines. This is exemplified by the Epi-Pluri-Score analysis (Lenz et al., 2015), and methylation in the *POU5F1* promoter (Fig. 4b; Supplemental file 1: Fig. S7a,b). We have previously described that age-associated DNAm patterns are reversed during reprogramming into iPSCs (Weidner et al., 2014) and all of our iPSC, iMSC, and iHPC lines maintained this rejuvenated epigenetic phenotype (Supplemental file 1: Fig. S7c).

**Fig. 4:**
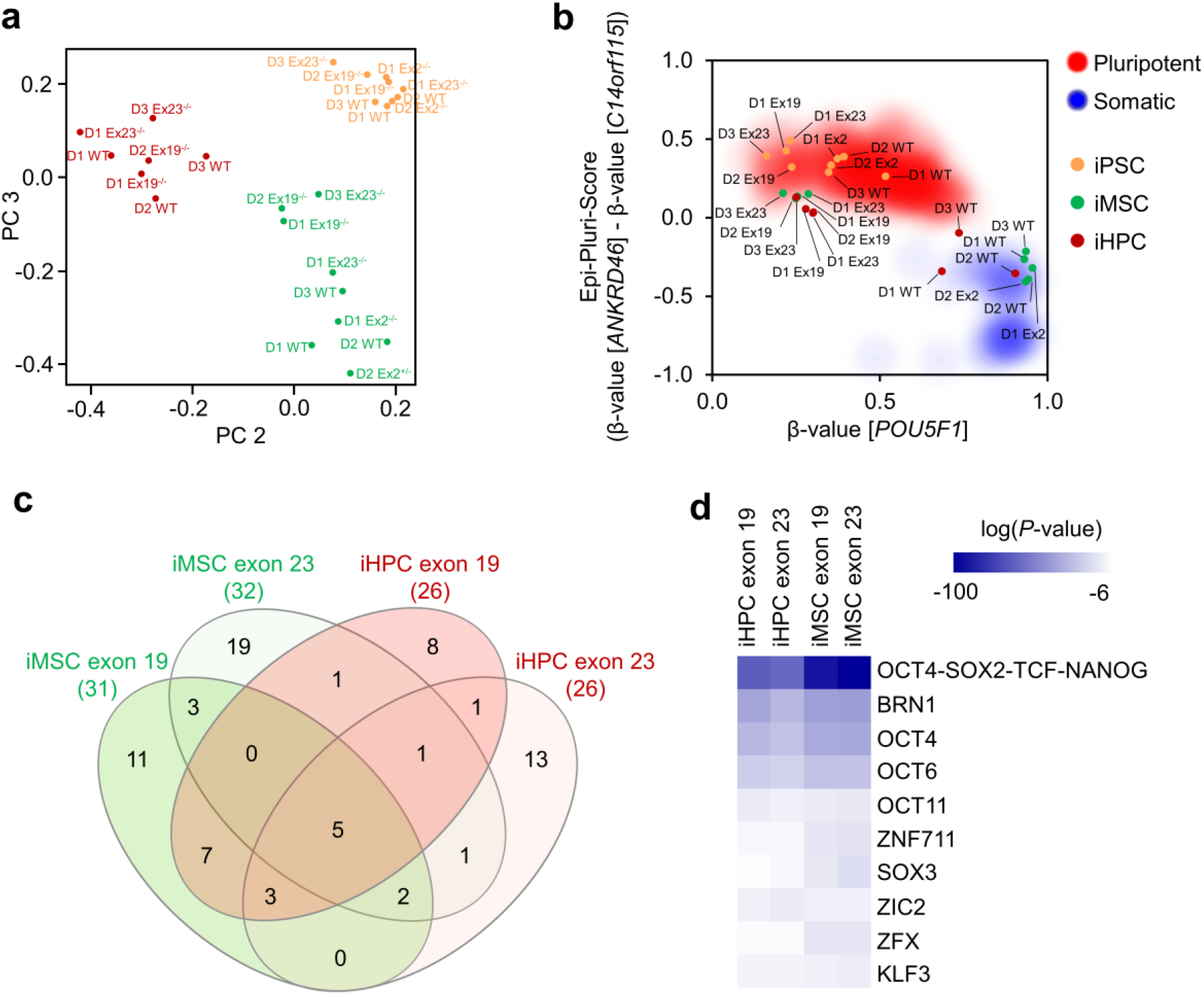
Comparison of DNA methylation profiles of all iPSCs, iMSCs and iHPCs. **a)** Principal component analysis of global DNA methylation profiles of all iPSC lines and derived iMSCs or iHPCs (D = donor, WT = wildtype, Ex = exon). **b)** Epi-Pluri-Score analysis of all generated cell lines and their progeny. **c)** Venn diagram showing overlap of CpGs that were >50% hypermethylated during differentiation of iPSCs into iMSCs or iHPCs (mean of two biological replica, each). **d)** Heatmap showing the top 10 transcription factor motifs enriched in these hypermethylated CpGs (compared to corresponding iPSCs).

Recently, it has been reported that genomic regions with low DNA methylation, termed long DNA methylation canyons, can form large loops connecting anchor loci of inter-chromosomal interaction (Zhang et al., 2020). *Dnmt3a* knockout mice demonstrated erosion of methylation canyons near developmental regulatory genes, like *HOX* clusters, or in association with HSC self-renewal genes (Jeong *et al*., 2018). Such analysis of large hypomethylated regions is better suited for whole genome bisulfite sequencing data, but we could also detect these canyons in our EPIC BeadChip data. In analogy to the previous studies in mice, particularly the canyon borders became hypomethylated in exon 19^-/-^ and exon 23^-/-^ knockout lines, as exemplarily depicted for the canyons in *PAX6* and *GATA2* (Supplemental file 1: Fig. S7d).

Finally, we focused on those few CpGs that still got highly methylated even without DNMT3A activity. These CpGs revealed a remarkable overlap for iMSCs and iHPCs, and for exon 19^-/-^ and exon 23^-/-^ (Fig. 4c). Most of these CpGs were located in enhancer regions. When we performed transcription factor binding motif analysis within a 100 bp window around the hypermethylated CpG sites, there was significant enrichment of pluripotency and development associated transcription factors, indicating that pluripotency-associated regulatory mechanisms are involved in the residual *de novo* DNAm during differentiation toward iMSCs or iHPCs (Fig. 4d).

### DNAm associated with *DNMT3A* mutations in AML is recapitulated in knockout iHPCs

To better understand if DNAm changes of our *DNMT3A* knockout iPSC clones are related to aberrant DNAm patterns in AML patients with *DNMT3A* mutations, we utilized 134 DNAm datasets of AML patients from The Cancer Genome Atlas (TCGA) without mutation in *DNMT3A* (n = 101), with R882 mutations (n = 18), or with other *DNMT3A* mutations (n = 15). AML patients with the R882 mutations revealed global hypomethylation, as compared to AML patients without *DNMT3A* mutation (Fig. 5a): 7 976 CpGs were more than 20% hypomethylated in R882 patients, whereas only 10 CpGs were 20% higher methylated. Notably, these hypomethylated CpGs showed on average also lower DNAm in our exon 19^-/-^ or exon 23^-/-^ iHPCs *versus* wildtype iHPCs (*P* < 1*10^−5^ as compared to all other CpGs for both comparisons; Fig. 5b,c). When we compared DNAm patterns in AML patients with other *DNMT3A* mutations to AML patients without *DNMT3A* mutations, the general hypomethylation was much less pronounced and only 896 CpGs revealed 20% lower DNAm with *DNMT3A* mutation, while 181 CpGs had at least 20% higher DNAm levels (Fig. 5d). Again, these hypomethylated CpGs were also hypomethylated in in our exon 19^-/-^ or exon 23^-/-^ iHPCs *versus* wildtype iHPCs (*P* < 1*10^−5^ for both comparisons, Fig. 5e,f). Thus, the DNAm patterns in AML patients that are related to *DNMT3A* mutations (R882 as well as other mutations) are partly recapitulated by our iPSC knockout models.

**Fig. 5:**
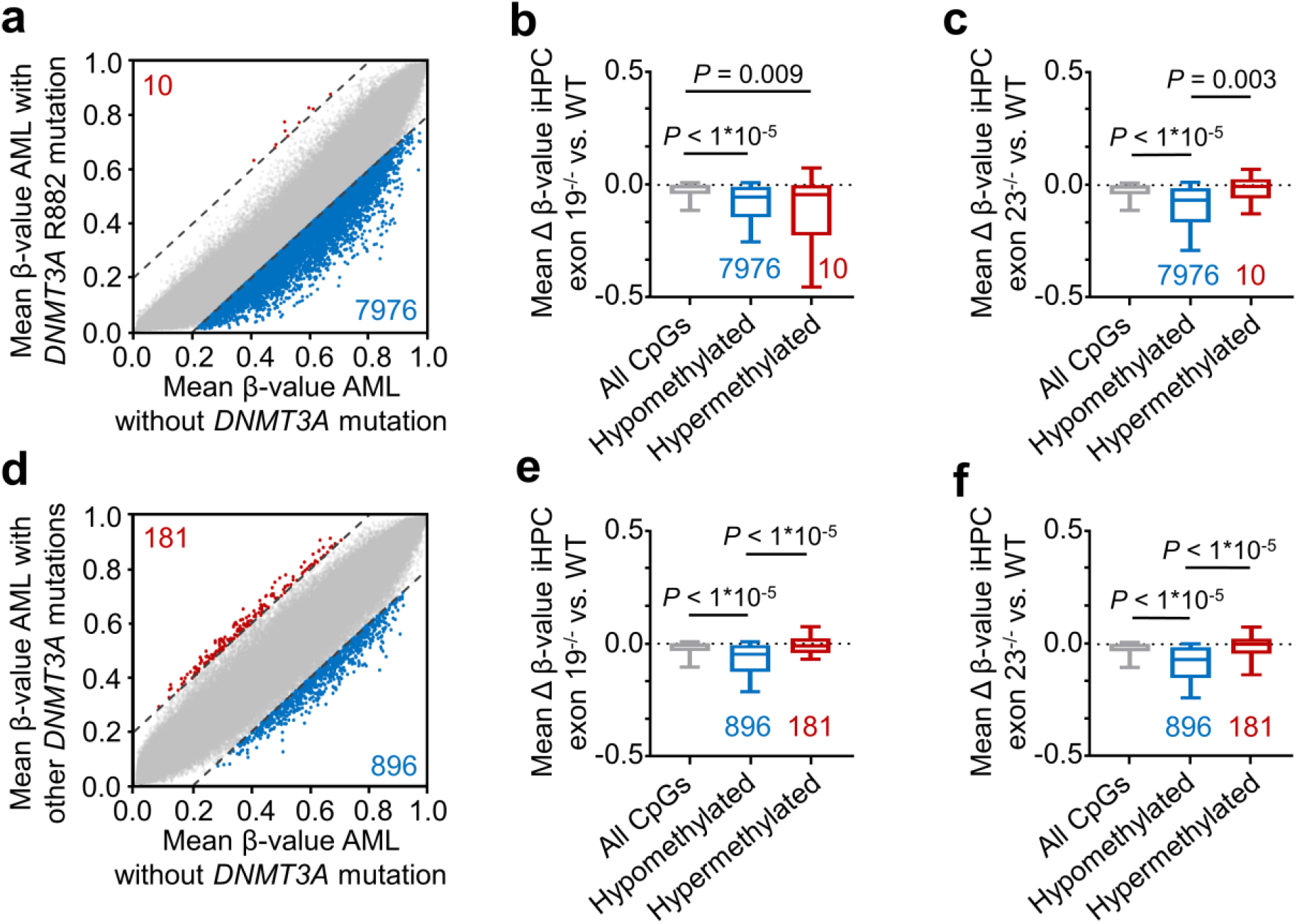
DNA methylation associated with *DNMT3A* mutations in acute myeloid leukemia. **a)** Scatter plot of DNAm levels in acute myeloid leukemia (AML) with *DNMT3A* R882 mutation (n = 18) plotted against data of patients without *DNMT3A* mutations (n = 101). The numbers of CpGs beyond the 20% threshold of differential DNAm are depicted. **b)** Box-whiskers plot (upper and lower quartiles, center line: median, whiskers: 10-90 percentile) showing mean hypo- and hypermethylation of the selected CpGs in (a) in the comparison *DNMT3A* exon 19^-/-^ *versus* wildtype (WT) iHPCs (mean of two biological replica). **c)** Box-whiskers plot of mean hypo- and hypermethylation of the selected CpGs in (a) in the comparison of *DNMT3A* exon 23^-/-^ *versus* WT. **d)** Scatter plot of DNAm levels in AML with *DNMT3A* mutations other than R882 (n = 15) plotted against data of patients without *DNMT3A* mutations. **e**,**f)** Box-whiskers plots of differentially methylated CpGs in AML with non-R882 mutations in *DNMT3A* in analogy to (b,c). All statistics were performed with 1-way ANOVA and Tukey’s PostHoc test.

### Exon 23 knockout shows growth advantage during competitive hematopoietic differentiation

We next wanted to better understand subclonal development during hematopoietic differentiation of iPSCs with and without *DNMT3A* knockouts. To this end, we transduced wildtype, exon 19^-/-^, or exon 23^-/-^ clones with lentiviral vectors containing unique molecular identifiers and the fluorophores Venus, Cerulean, or mCherry (Fig. 6a). The three syngeneic iPSC lines were then mixed with equal cell numbers and seeded on the microcontact-printed plates for EB formation (Fig. 6b). After hematopoietic differentiation, flow cytometric analysis demonstrated growth advantage of mCherry labeled exon 23^-/-^ clones (Fig. 6c), which is in line with the higher iHPC numbers (Fig. 3c). Furthermore, we analyzed the fluorophore-specific genetic barcodes by amplicon deep-sequencing: again exon 23^-/-^ cells had clear growth advantage with increasing fractions upon EB formation, iHPC production, and after additional long-term culture expansion for 28 days, whereas counts for exon 19^-/-^ cells decreased with time (Fig. 3d). Subsequently, the fractions of corresponding unique molecular identifier were tracked over the different time points demonstrating that hematopoietic differentiation of iPSCs is a multiclonal event (Fig. 3e). While some exon 23^-/-^ barcodes made up to 1.51% of all reads there was no evidence for a clear oligo clonal composition after long-term expansion.

**Fig. 6:**
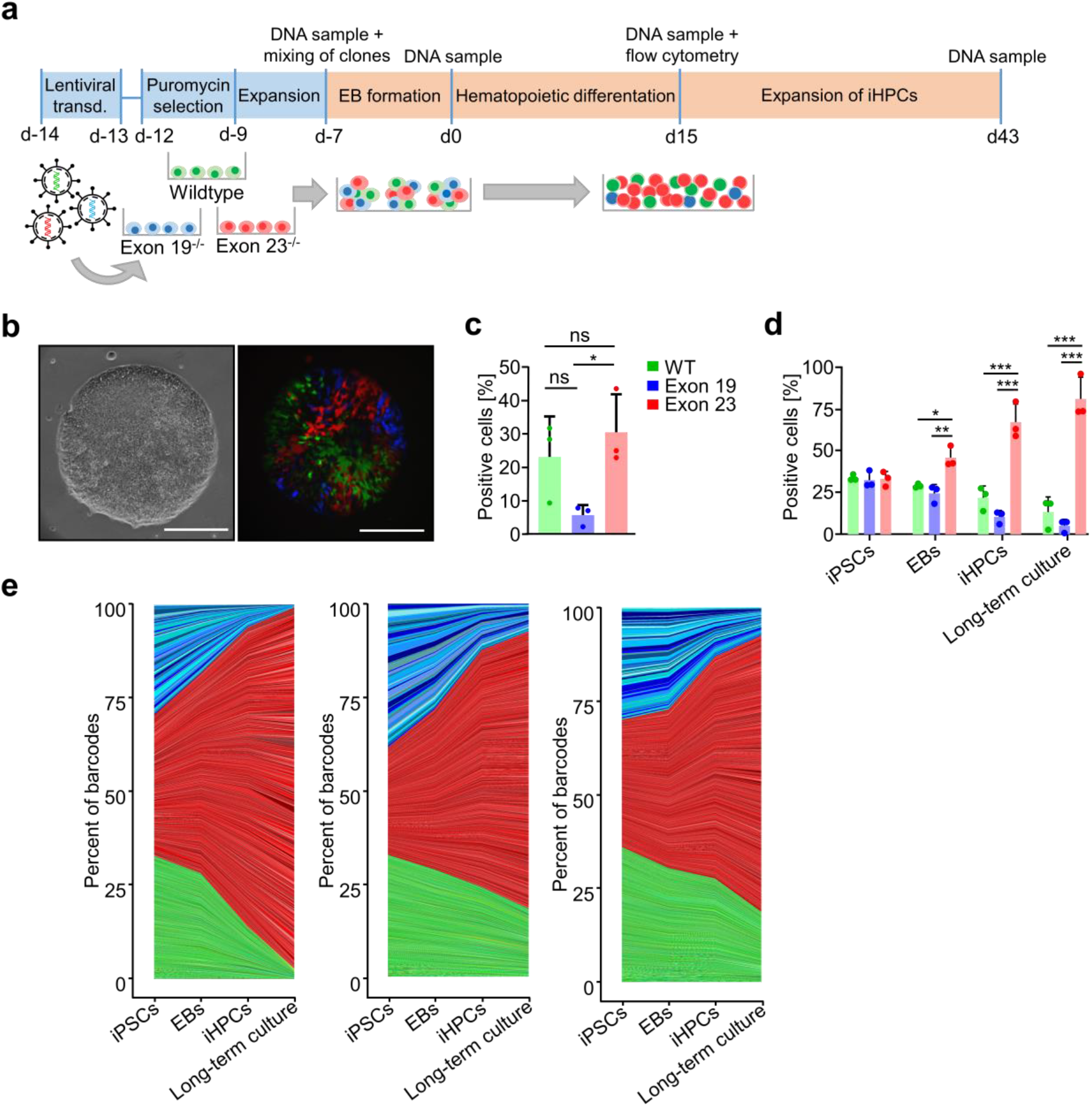
Competitive growth advantage of exon 23^-/-^ cells during hematopoietic differentiation. **a)** Syngeneic wildtype, exon 19^-/-^ and exon 23^-/-^ iPSCs were transduced with a lentiviral barcoding system and the fluorophores Venus, Cerulean, and mCherry, respectively. The schematic representation depicts competitive growth in co-culture during embryoid body (EB) formation, hematopoietic differentiation, and additional long-term culture (n = 3, all donor 1). **b)** Representative phase contrast and fluorescence microscopic images of EBs with mixed populations at day -4 (day 3 of EB growth). **c)** Flow cytometry analysis of iHPCs after 15 days of hematopoietic differentiation (only cells with a clear fluorescent signal were considered). n = 3 (mean ± SD). **d)** Alternatively, we analyzed the fluorophore-specific genetic barcodes by amplicon sequencing after mixing of the iPSCs (d-7), in EBs (d0), in iHPCs (d15), and after additional long-term expansion (d43). **e)** Area plots show clonal growth during differentiation and expansion of the three replicates. The graded colors depict corresponding unique molecular identifiers of the lentiviral barcoding. All statistics were performed with 1-way ANOVA and Tukey’s PostHoc test. * *P* < 0.05, ** *P* < 0.01, *** *P* < 0.001.

## Discussion

So far, the sequel of losing DNMT3A activity for hematopoietic development has mostly been investigated in the murine system by knocking out exon 19 or exon 18-20 of *Dnmt3a* with the Cre/loxP system (Challen et al., 2011; Izzo et al., 2020), or by targeting exon 19 with CRISPR/Cas9 (Yagi et al., 2020). Two groups established *DNMT3A* knockouts in human ESCs and iPSCs by targeting exon 19 (Liao *et al*., 2015; Madsen *et al*., 2020), but they did not describe their hematopoietic differentiation potential. iPSCs are a valuable model system for hematopoietic differentiation, because they facilitate systematic and clonal analysis in homogenous cell preparations (Shi et al., 2017).

Our results demonstrate that iPSCs with knockout of *DNMT3A* remain capable of differentiating toward hematopoietic lineages. Despite the pronounced differences in DNAm profiles of wildtype and *DNMT3A*^-/-^ clones, we did not find any significant differences in gene expression, indicating that *de novo* DNA methylation is not essential for differentiation into iHPCs. It needs to be considered that DNAm patterns of wildtype and knockout clone-derived iPSCs, iHPCs, and iMSCs still clustered together, probably due to cell-type specific hypomethylation, which may be functionally more relevant. Furthermore, other epigenetic modifications, such as histone modifications and chromatin conformation, may be primary drivers of directed cellular differentiation.

There are many different site-specific *DNMT3A* mutations in leukemia and CHIP (Brunetti *et al*., 2017; Ley *et al*., 2010) and therefore, it might be anticipated that targeting of specific exons is of functional relevance. Exon 2 mutations in AML have been described before but its functional impact remains unclear and this exon is not associated with a functional protein domain (Lu et al., 2013; Patterson et al., 2016). Mutations in exon 19 are highly abundant and may result in a functional loss (Holz-Schietinger et al., 2012; Sandoval et al., 2019). Exon 19 plays a role for DNA binding and modulation of the methyltransferase domain (Zhang et al., 2018). More research has been performed for mutations in exon 23, particularly the hotspot mutation R882, which often results in loss of DNAm (Emperle *et al*., 2019; Holz-Schietinger *et al*., 2012). This region is also involved in the methyltransferase domain and it is essential for interacting with the DNA backbone during DNMT3A-DNMT3A tetramer formation (Zhang *et al*., 2018). Our results support the notion that there may be functional differences between the different *DNMT3A* knockouts, but it needs to be considered that for each exon only two knockout lines have been generated and there is notorious variation in the differentiation potential of such cell lines. Furthermore, there were hardly differences in the DNA methylation profiles of iHPCs with exon 19^-/-^ and exon 23^-/-^. Additional research is therefore necessary to ultimately answer the question how aberrant splicing and specific mutations affect hematopoietic differentiation.

Only very few CpGs consistently gained DNAm in our exon 19^-/-^ and exon 23^-/-^ clones, and these were associated with pluripotency and development associated transcription factor binding sites. In primary hematopoiesis, Dnmt3a has been shown to be a critical participant in the epigenetic silencing of HSC regulatory genes, thereby enabling efficient differentiation (Challen *et al*., 2011). In contrast, Dnmt3b is also highly abundant in HSCs, but rather in a catalytically inactivated form with just some residual activity (Challen et al., 2014; Okano *et al*., 1999). The *de novo* methylation of these few CpGs might therefore be attributed to DNMT3B.

Acute myeloid leukemia is associated with characteristic aberrations in the DNA methylation pattern (Bozic et al., 2021). Patient samples with the hotspot mutation R882 revealed prominent hypomethylation compared to other *DNMT3A* mutations, as described before (Russler-Germain *et al*., 2014). Our results demonstrate that iPSC-derived model iHPCs can recapitulate some epigenetic effects of R882 mutated, and even in non-R882 mutated AML. Thus, aspects of AML-related DNAm patterns can be modelled by our system, which may be useful for further mechanistic analysis.

Lastly, we investigated if our iPSC model system also reveals subclonal dominance of *DNMT3A* knockouts in a competitive differentiation assay with long-term culture. In fact, exon 23^-/-^ iHPCs revealed growth advantage over wildtype and exon 19^-/-^ lines but there was no evidence for dominant subclones. In contrast, the lentiviral barcoding system did revealed dominant subclones in iMSCs, even without modifications in *DNMT3A* (Hollmann et al., 2020). It is conceivable that differentiation of iHPCs is more homogeneous than iMSCs and that the effects of *DNMT3A* knockouts become only apparent after even longer culturing periods. Either way, a moderate growth advantage of clones with specific aberrations in *DNMT3A* might be a functional correlate to malignant transformation.

Site specific DNAm is an important epigenetic mechanism during differentiation. It is therefore remarkable that there is hardly any impact on lineage specific differentiation of our iPSC clones with homozygous knockout of exon 19 or 23, despite severe interference with *de novo* methylation. To better understand the functional relevance, it will therefore be important to gain better insight into cell-type specific hypomethylation and the interplay with other epigenetic modifications. The mechanisms that may result in the functional differences between *DNMT3A* knockout lines remain to be further elucidated, also with regard to patient-specific iPSC lines. Our model system can help to get a better understanding on the relevance of DNMT3A during hematopoietic development and malignant transformation.

## Experimental procedures

### iPSC lines, ethics approval, and consent to participate

iPSCs of donor 1 and donor 2 were reprogrammed from MSC preparations with episomal plasmids and characterized, as described before (Goetzke et al., 2018). iPSCs of donor 3 were generated from KIT+ cells from bone marrow aspirates of a mastocytosis patient, with the CytoTune-iPS Sendai Reprogramming Kit (Thermo Fisher Scientific, Waltham, MA, USA). This line was also carefully characterized and revealed a *KRAS* mutation A146T. All samples were taken after informed and written consent and the study was approved by the Ethic Committee of the Use of Human Subjects at the University of Aachen (permit numbers: EK128/09 and EK206/09, respectively). iPSCs were cultured on tissue culture plastic coated either with vitronectin (0.5 mg/cm^2^; Stemcell Technologies, Vancouver, Canada) or Matrigel Matrix (Corning, Corning, NY, USA) in StemMACS iPS-Brew XF (Miltenyi Biotec, Bergisch Gladbach, Germany) with 100 U/mL penicillin and 100 μg/mL streptomycin. Pluripotency was validated by Epi-Pluri-Score (Cygenia GmbH, Aachen, Germany) (Lenz *et al*., 2015).

### CRISPR/Cas9n knockout of *DNMT3A*

Knockouts of exon 2, 19, and 23 of *DNMT3A* were generated with CRISPR/Cas9n double nicking approach (Ran et al., 2013). Two pairs of gRNAs were designed with the help of the CRISPOR tool (Concordet and Haeussler, 2018) (Supplemental file 1: Fig. S1a & Tab. S1). gRNA primers were individually ligated into a variant of the pX335 vector (Addgene #42335) additionally containing the reporter protein GFP and a puromycin selection cassette (Cong et al., 2013; Zhang et al., 2014). Transfection of 10^6^ iPSCs with the four gRNA plasmids (2.5 µg each) was performed with the NEON transfection system kit (Thermo Fisher Scientific) and after 1 day transfected cells were selected with 0.4 µg/mL puromycin for 48 h. Single colonies were picked and screened for *DNMT3A* deletions by PCR and Sanger sequencing (primers: Supplemental file 1: Tab. S2). Knockout was further confirmed with semi-quantitative real-time PCR and Western blots, pluripotency was tested with immunofluorescence and trilineage differentiation assays as described in Supplemental file 1.

### Differentiation of iPSCs into the mesenchymal lineage

For iMSC generation, iPSC medium was switched to MSC medium, consisting of Dulbecco’s modified Eagle’s medium-low glucose (Thermo Fisher Scientific) supplemented with 10% human platelet lysate, and 5 U/ml heparin (Rotexmedica, Trittau, Germany) for 35 days, as described before (Frobel et al., 2014). Surface marker expression was measured with flow cytometry (Supplemental file 1).

### Hematopoietic progenitor differentiation

To differentiate iPSCs into hematopoietic progenitors, we adjusted the protocol from Liu et al. (Liu et al., 2015): EBs were generated from iPSCs by self-detachment from microcontact-printed vitronectin spots generated with PDMS stamps (Elsafi Mabrouk et al., 2020). EBs were carefully resuspended in serum free medium containing 50% IMDM, 50% Ham’s F12, 0.5% BSA, 1% chemically defined lipid concentrate, 2 mM GlutaMAX (all Thermo Fisher Scientific), 400 µM 1-thioglycerol, 50 µg/mL L-ascorbic acid, and 6 µg/mL holo transferrin (all Sigma Aldrich, St. Louis, MO, USA) supplemented with 10 ng/mL FGF-2 (Peprotech, Hamburg, Germany), 10 ng/mL BMP-4 (Miltenyi Biotec), and 10 µM Y-27632 (Abcam, Cambridge, Great Britain). EBs were seeded on 0.1% gelatin coated 6-well plates. From day 2 to 16 EBs were cultured in serum free medium supplemented with 10 ng/mL FGF 2, 10 ng/mL BMP-4, 50 ng/mL SCF, 10 ng/mL VEGF-A (all Peprotech) and 10 U/mL penicillin/streptomycin (Thermo Fisher Scientific). The non-adherent iHPCs were harvested at day 16 and separated with a 40 µm cell strainer. Immunophenotype was tested with flow cytometry and stem cell potential with colony forming unit assays as described in Supplemental file 1.

### DNA methylation analysis

Genomic DNA was isolated with the NucleoSpin Tissue kit (Macherey Nagel) and subsequently bisulfite converted and hybridized to the Infinium MethylationEPIC BeadChip (Illumina, San Diego, California, USA) at Life and Brain GmbH (Bonn, Germany). Data was preprocessed with the Bioconductor Illumina Minfi package for R (Aryee et al., 2014; Fortin et al., 2017) and normalized with ssnoob. CpGs on X and Y chromosomes, SNPs and cross-reactive sites were excluded, resulting in 741 896 CpGs. For comparison with other datasets (iHPC: GSE60811 and GSE119079; whole blood, granulocytes, B cells, CD4+ T cells, CD8+ T cells, NK cells, monocytes: all GSE35069; CD34+ cells isolated from human cord blood: GSE40799; primary MSCs: GSE113527; AML data obtained from The Cancer Genome Atlas) we further focused on CpGs that were presented by the Illumina Infinium HumanMethylation450 BeadChip and the Infinium MethylationEPIC BeadChip. Principal component analysis and multidimensional scaling plots were prepared in R using the ggplot2 package version 3.3.3 (Wickham, 2011). Heatmaps were generated with the MultiExperiment Viewer (MeV; version 4.9.0). Gene ontology analysis was performed on genes with differentially methylated CpGs in the promoter region (located in TSS1500, TSS200, and 5′UTR) in R with the missMethyl package (Phipson et al., 2016). Categories comprising more than 1 000 genes were not considered and similar categories are only listed once. Venn diagrams were prepared with the InteractiVenn tool (Heberle et al., 2015). Motif analysis was performed with the Hypergeometric Optimization of Motif EnRichment (HOMER) suite (Heinz et al., 2010) with a 100 bp frame around hypermethylated CpGs. Canyons were defined as 30 neighboring CpG sites within 3 kb and an average β-value below 0.15. Epigenetic age was predicted using Horvath’s skin and blood clock (Horvath et al., 2018).

### Gene expression analysis

For transcriptomic analysis, total RNA was isolated from iHPCs at d16 of differentiation with the NucleoSpin RNA Plus kit (Macherey Nagel). mRNA was prepared with the QuantSeq 3’ mRNA-Seq Library Prep Kit FWD for Illumina (Lexogen, Vienna, Austria) and sequenced with 10 million 50 bp single-end reads on the HiSeq 2500v4 (Illumina) at Life and Brain GmbH. Raw data underwent quality control with Trim Galore and sequences were aligned to the human genome (hg38) by bowtie2 (Langmead and Salzberg, 2012) aligner with default parameters. Differential expression analysis was done by DESeq2 (Love et al., 2014) with the adjusted p-value threshold of 0.05. Scatter plots were generated with ggplot2 (Wickham, 2011) in R.

### Genetic barcoding of iPSC clones with RGB LeGO vectors

Generation of barcoded RGB vectors was performed as described in detail before (Hollmann *et* al., 2020). LeGO vectors expressing either the fluorophores Venus, Cerulean, or mCherry (pRRL-PPT-CBX3-EFS-Cerulean-P2A-Puro, pRRL-PPT-CBX3-EFS-mCherry-P2A-Puro, pRRL-PPT-CBX3-EFS-Venus-P2A-Puro (Selich et al., 2019)) were ligated with unique molecular identifiers containing random as well as color-specific nucleotide sequences (Selich et al., 2016) (RGB-vectors; kindly provided by the Medizinische Hochschule Hannover, Germany). Wildtype, exon 19^-/-^ and exon 23^-/-^ iPSCs were transduced with the Venus, Cerulean, or mCherry libraries, respectively, and used for a competitive hematopoietic differentiation assay as described in detail in Supplemental file 1.

### MiSeq analysis of barcoded cells

MiSeq was performed as described in detail before (Hollmann *et al*., 2020). In brief, DNA was isolated with the NucleoSpin Tissue kit (Macherey Nagel), barcodes were amplified, tagged with Illumina sample barcodes, samples were pooled equimolarly, and a 4 nM library was prepared for sequencing with a 50% spike in of PhiX Control DNA and the MiSeq Reagent Kit v2 (all from Illumina). Libraries were sequenced in 250 bp paired-end mode on an Illumina MiSeq Benchtop Sequencer. Raw data was analyzed with Python. Unique molecular identifiers of the RGB-vectors were extracted from FastQ files and grouped according to color-specific sequences resembling the different cell lines. Two mismatches were allowed for grouping of these color-specific identifiers. Finally, directional clustering from UMI-tools (Smith et al., 2017) was used to specify unique molecular identifiers for each color-group. Area plots were prepared with ggplot2 in R.

### Statistical analysis

Data is provided with standard deviation (SD). Significance of flow cytometry data was calculated by 2-way ANOVA with Tukey’s PostHoc test. 1-way ANOVA with Tukey’s PostHoc test was used for iHPC counts, CFU colonies, for comparison of methylation data with AML patients, and for flow cytometry data of RGB cells (GraphPad version 9.1.1.). Significantly differentially expressed genes were tested in R with DESeq2 using the default Wald test.

## Acknowledgements

This research was supported by the German Research Foundation (DFG: WA 1706/8-1, WA 1706/12-1); the Interdisciplinary Center for Clinical Research (IZKF) within the faculty of Medicine at RWTH Aachen University (O3-3); Deutsche Krebshilfe (TRACK-AML); the Federal Ministry of Education and Research (BMBF: VIP + Epi-Blood-Count); and by the Flow Cytometry Facility, a core facility of the IZKF Aachen. Furthermore, we thank Janik Boehnke, Kathrin Olschok and Marcelo Szymanski de Toledo for discussion and advice on iPS cell differentiation towards hematopoietic cells.

## Author contributions

O.C. performed experiments and analyzed data. J.Fra. supported DNAm and RGB barcode analysis. J.Fro. created exon 23 knockout lines. P.G. performed RGB barcoding experiments. C.C.K. supported RNAseq analysis and motif analysis. S.S. performed trilineage differentiation of iPSCs and helped with hematopoietic differentiation. S.N. supported iMSC differentiation. M.Z. contributed to study design. O.C. and W.W. designed the study, wrote the first draft of the manuscript and all authors approved the final manuscript.

## Supplemental files

**Supplemental file 1. Combined PDF of Supplemental figures S1 – S7, Supplemental tables S1 – S4 and Supplemental experimental procedures**.

Fig. S1: Generation of CRISPR/Cas9n knockout iPSC clones. Fig. S2: Pluripotency and global DNA methylation in iPSC lines. Fig. S3: Gene ontology analysis and overlap of differentially methylated CpGs upon mesenchymal differentiation. Fig. S4: Characterization of iPSC-derived hematopoietic progenitor cells. Fig. S5: DNA methylation analysis of hematopoietic progenitor cells. Fig. S6: Comparison of iHPCs with exon 19 or exon 23 knockout. Fig. S7: Gene methylation, age prediction and canyon analysis of iPSCs, iMSCs and iHPCs. Tab. S1: gRNAs for CRISPR. Tab. S2: Flanking primers for PCR. Tab. S3: Exon specific primers for qPCR. Tab. S4: qPCR primers for the trilineage assay.

**Supplemental file 2. Tab. S5: Differentially methylated CpGs in iPSC WT *versus DNMT3A* knockouts**.

CpG sites that are either 50% hypo- or hypermethylated in *DNMT3A* exon 2^-/-^, exon 19^-/-^, or exon 23^-/-^ iPSCs compared to wildtype iPSCs with associated genes, gene groups, relation to CpG islands, chromosomes, positions, mean beta-values of the knockout and wildtype cells, and the difference in methylation between knockout and wildtype.

**Supplemental file 3. Tab. S6: Differentially methylated CpGs in iPSCs *versus* iMSCs (WT or *DNMT3A* knockouts)**.

CpG sites that are either 50% hypo- or hypermethylated in *DNMT3A* wildtype, exon 2^-/-^, exon 19^-/-^, or exon 23^-/-^ iMSCs compared to their iPSC counterparts with associated genes, gene groups, relation to CpG islands, chromosomes, positions, mean beta-values of the knockout and wildtype cells, and the difference in methylation between knockout and wildtype.

**Supplemental file 4. Tab. S7: Differentially methylated CpGs in iPSCs *versus* iHPCs (WT or *DNMT3A* knockouts)**.

CpG sites that are either 50% hypo- or hypermethylated in *DNMT3A* wildtype, exon 19^-/-^, or exon 23^-/-^ iHPCs compared to their iPSC counterparts with associated genes, gene groups, relation to CpG islands, chromosomes, positions, mean beta-values of the knockout and wildtype cells, and the difference in methylation between knockout and wildtype.

**Supplemental file 5. Tab. S8: Differentially methylated CpGs in iHPC WT *versus DNMT3A* knockouts**.

CpG sites that are either 50% hypo- or hypermethylated in *DNMT3A* exon 19^-/-^, or exon 23^-/-^ iHPCs compared to wildtype iHPCs with associated genes, gene groups, relation to CpG islands, chromosomes, positions, mean beta-values of the knockout and wildtype cells, and the difference in methylation between knockout and wildtype.

**Supplemental file 6. Tab. S9: RNA-seq data of iHPCs**.

Genes that are either four-fold downregulated or upregulated (log_2_ fold change <-2 or >2) in *DNMT3A* wildtype *versus* exon 19^-/-^, wildtype *versus* exon 23^-/-^, or exon 19^-/-^ *versus* exon 23^-/-^ with gene name, base mean, log_2_ fold change, *P*-value, adjusted *P*-value, and read counts per million of knockout and control cells.

