## Supplemental file 1 for "*DNMT3A* knockouts in human iPSCs prevent *de novo* DNA methylation and reveal growth advantage during hematopoietic differentiation"

### *DNMT3A* knockouts in human iPSCs prevent *de novo* DNA methylation and reveal growth advantage during hematopoietic differentiation

Olivia Cypris, Julia Franzen, Joana Frobel, Philipp Glück, Chao-Chung Kuo, Stephani Schmitz, Selina Nüchtern, Martin Zenke, and Wolfgang Wagner

### Supplemental figures

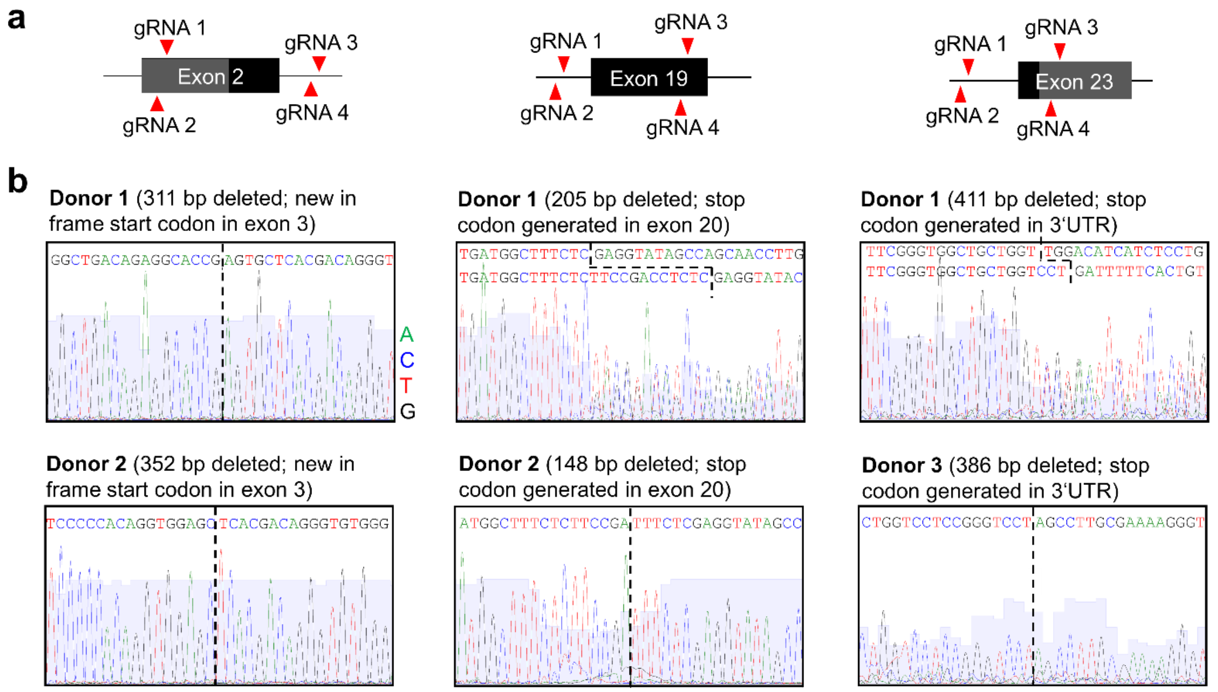


**Fig. S1: Generation of CRISPR/Cas9n knockout iPSC clones.**

**(a)** Schematic representation of gRNA design for exon 2 knockout, exon 19 knockout, and exon 23 knockout, respectively. The boundary between the grey and black box of exon 2 is the transcription start site. Red arrows depict cutting sites of guide RNAs (gRNAs). **(b)** Sanger sequencing results are depicted for the two clones for each exon, which were subsequently used in this study. Dotted lines = CRISPR cutting site. In donor 1 exon 19 and exon 23, each allele was cut at different sites as indicated.

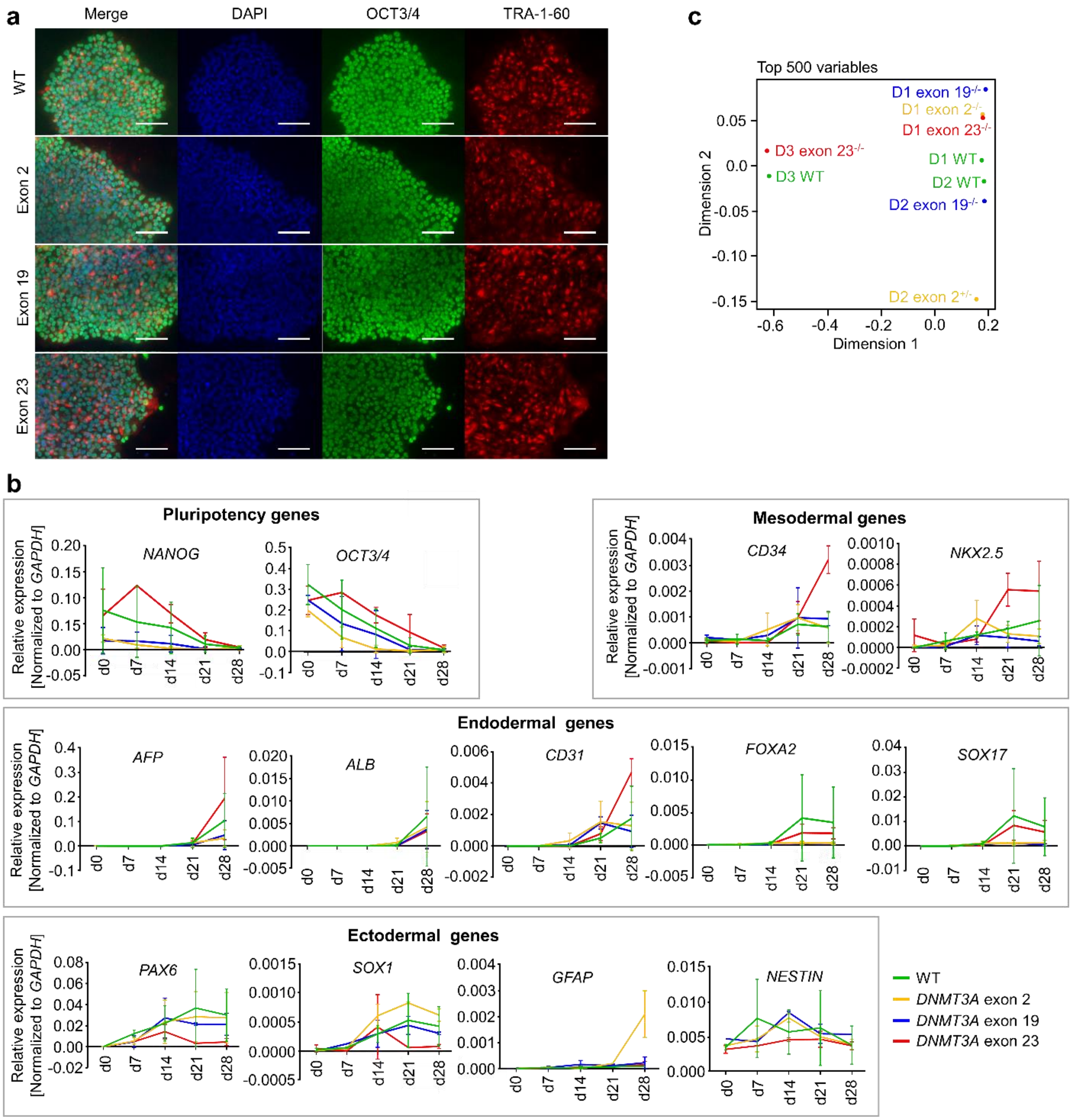


**Fig. S2: Pluripotency and global DNA methylation in iPSC lines.**

**(a)** Immunofluorescence staining of nuclei with DAPI (blue) and the pluripotency markers OCT3/4 (green) and TRA-1-60 (red) of representative clones of wildtype (WT) and knockout lines. **(b)** Relative expression of pluripotent, mesodermal, endodermal and ectodermal markers after trilineage differentiation of all iPSC lines as measured by quantitative real-time PCR. Expression values were normalized to the housekeeping gene *GAPDH*. n = 3 for WT, n = 2 for each knockout with duplicates for each measurement. Data is presented as mean ± SD. **(c)** Multidimensional scaling plot of all iPSC lines with the top 500 most variable CpG sites (D = donor).

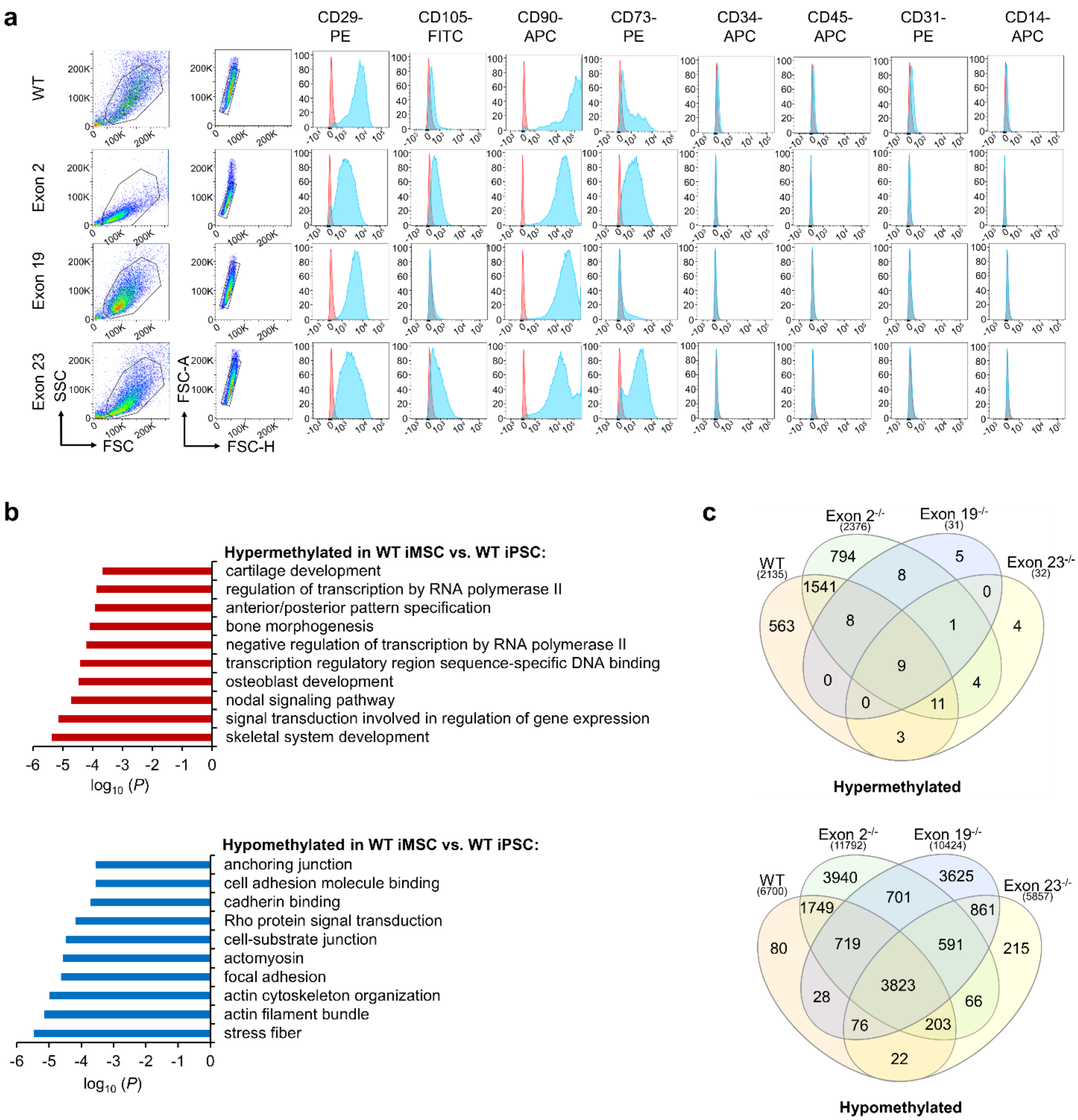


**Fig. S3: Gene ontology analysis and overlap of differentially methylated CpGs upon mesenchymal differentiation.**

**(a)** Flow cytometry analysis of iMSCs after 35 days of differentiation. WT = wildtype, FSC = forward scatter, SSC = side scatter. **(b)** Gene ontology analysis of genes with >50% differentially methylated CpG sites in the promoter region. **(c)** Venn diagram showing the overlap of 50% hyper- (left) and hypomethylated (right) CpG sites during differentiation of iPSCs to iMSCs, when comparing wildtype (WT) and *DNMT3A* knockout clones (means of two biological replica per comparison for exon $19^{-/-}$ and exon $23^{-/-}$).

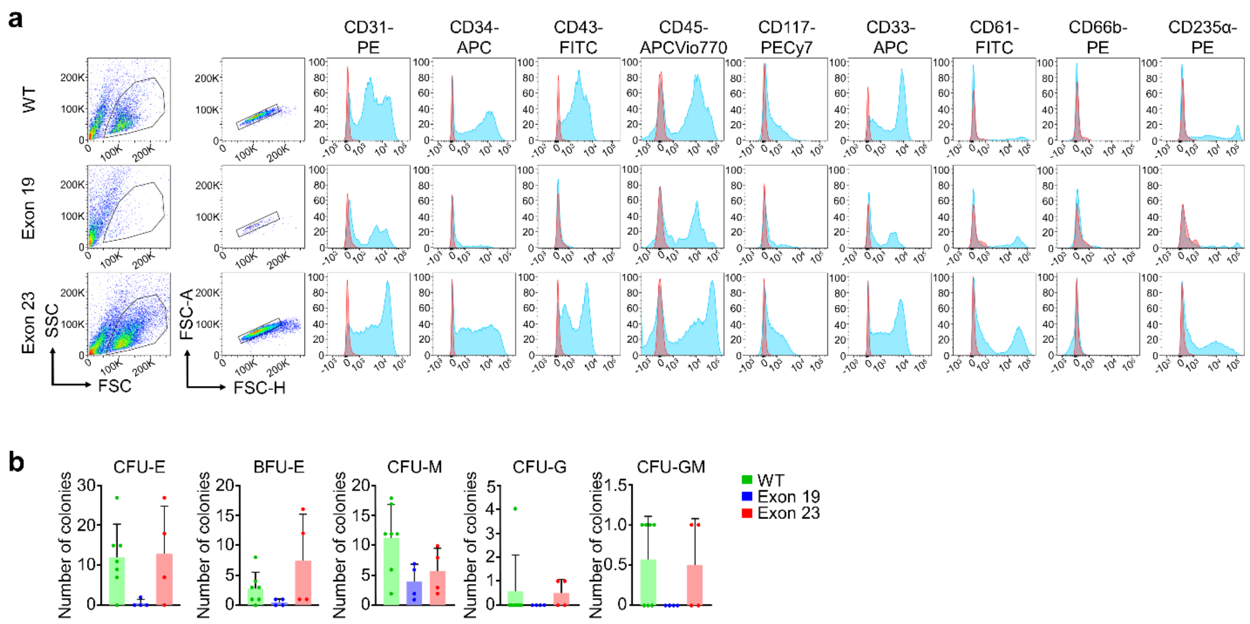


**Fig. S4: Characterization of iPSC-derived hematopoietic progenitor cells.**

**(a)** Flow cytometry analysis of iHPCs after 16 days of differentiation. WT = wildtype, FSC = forward scatter, SSC = side scatter. **(b)** Total numbers of colony forming units (CFUs) for each colony type. CFU-E = CFU erythrocyte CFU; BFU-E = burst forming unit erythrocyte; CFU-M = CFU macrophage; CFU-G = CFU granulocyte; CFU-GM = CFU granulocyte, macrophage; CFU-GEMM = granulocyte, erythrocyte, macrophage, megakaryocyte. Statistics were calculated with 1-way ANOVA and Tukey‘s PostHoc test. n = 7 for WT, n = 4 for each knockout with duplicates for each measurement. Data is presented as mean ± SD.

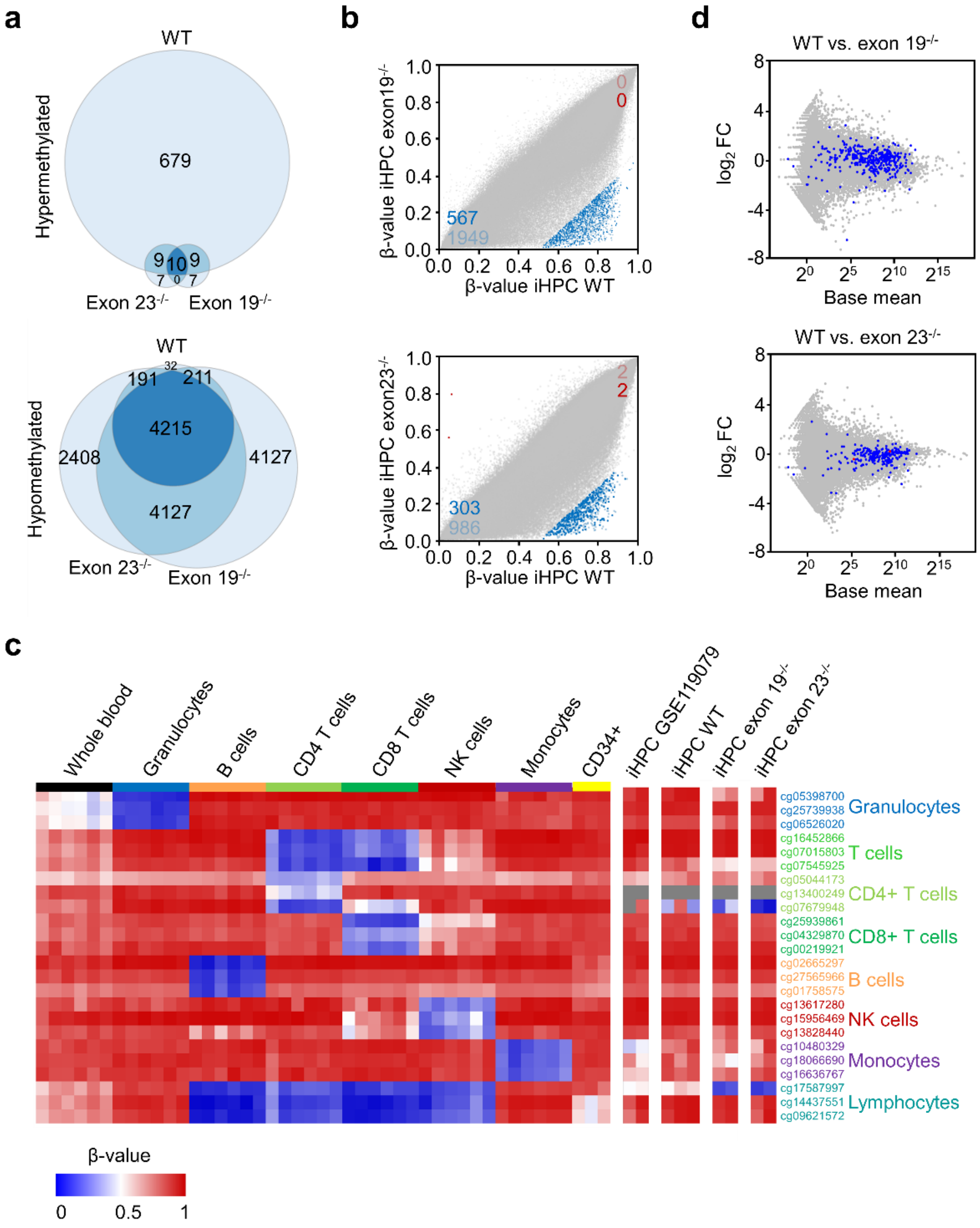


**Fig. S5: DNA methylation analysis of hematopoietic progenitor cells.**

**(a)** Venn diagram showing the overlap of CpG sites that become at least 50% hyper- or hypomethylated during differentiation of iPSCs into iHPCs (mean of two replica for each comparison). WT = wildtype. **(b)** Scatter plots showing β-values for all CpG sites measured in iHPC wildtypes and exon 19 or exon 23 knockouts in grey, 50% hypo- (blue) and 50% hypermethylated (red) CpG sites. The darker blue and red dots and numbers represent CpG sites in the promoter region. **(c)** Heatmap of DNA methylation levels at CpGs that are specifically hypomethylated in specific hematopoietic cell types. The profiles of whole blood, granulocytes, B cells, CD4+ T cells, CD8+ T cells, NK cells, monocytes were taken from GSE35069 (Reinius et al., 2012); CD34+ cells isolated from human cord blood from GSE40799 (Weidner et al., 2013); and DNAm profiles of iHPC from GSE119079 (Cypris et al., 2019) and our current study. Gray areas indicate missing data. **(d)** Differentially promoter-methylated CpG associated genes in iHPC wildtypes compared to exon $19^{-/-}$ or exon $23^{-/-}$ are plotted in blue (50% hypomethylated) or red (50% hypermethylated) in the graphs from Fig. 3h.

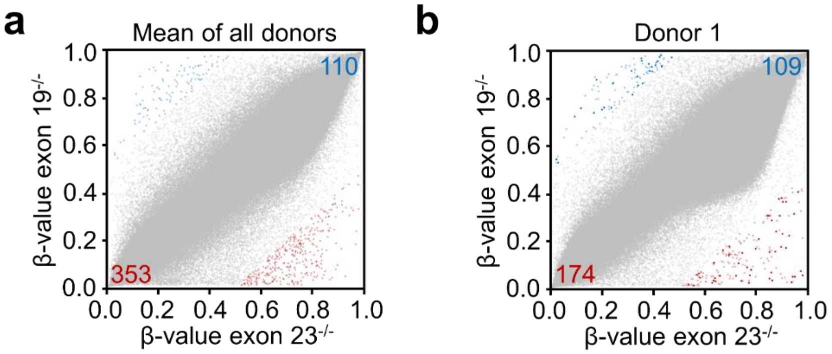


**Fig. S6: Comparison of iHPCs with exon 19 or exon 23 knockout.**

**(a)** Scatter plot showing mean β-values for all CpG sites measured in iHPC exon 19 and exon 23 knockout cells of all donors in grey with 50% hypo- and 50% hypermethylated CpG sites indicated in blue or red, respectively. **(b)** Same as (a) but showing values for the comparison of donor 1 clones only.

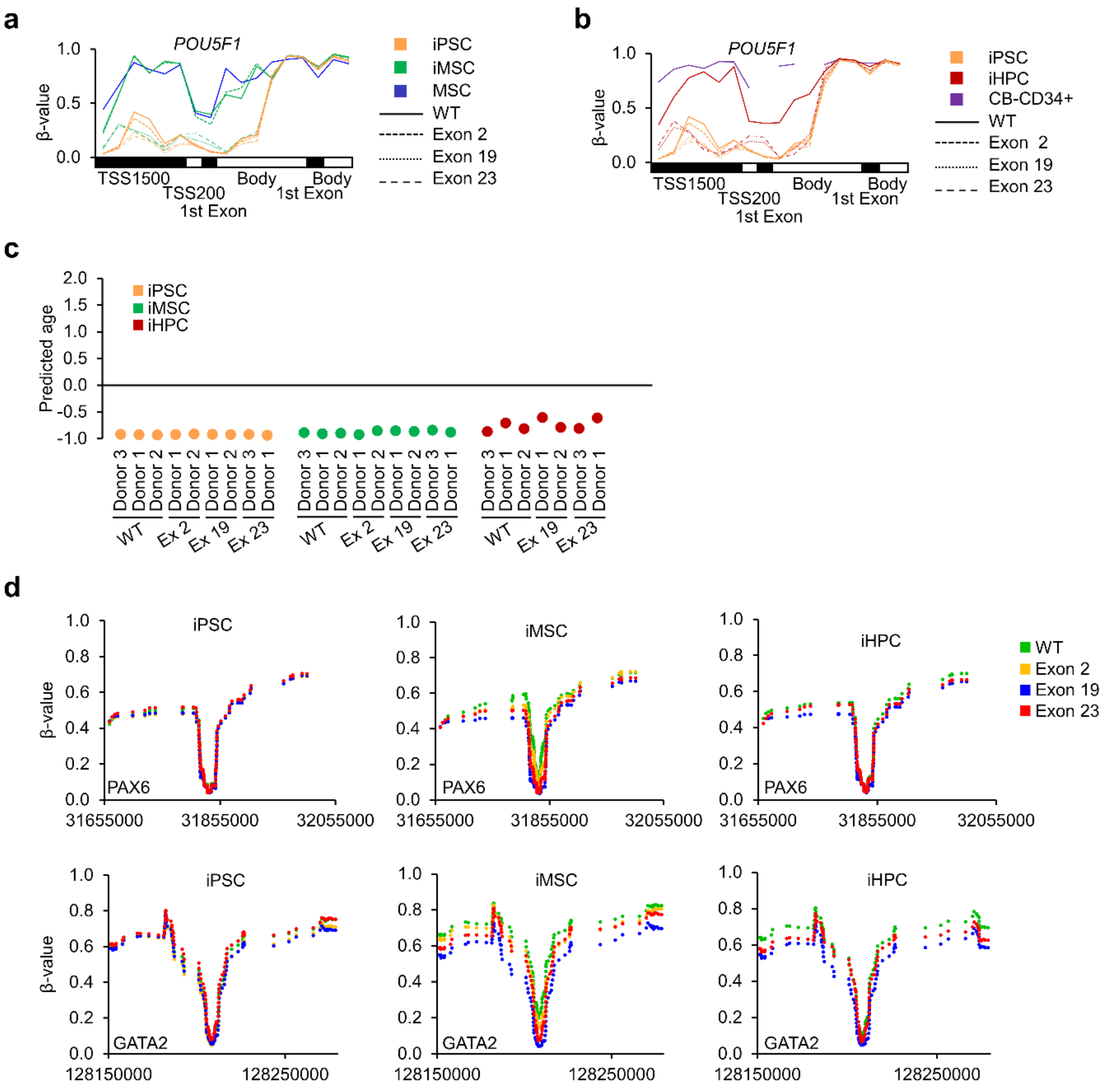


**Fig. S7: Gene methylation, age prediction and canyon analysis of iPSCs, iMSCs and iHPCs.**

**(a)** DNAm of CpGs within the gene of *POU5F1* in iPSCs, iMSCs, and primary MSCs (GSE113527) (De Witte et al., 2018). **(b)** DNAm of CpGs within the gene *POU5F1* in iPSCs, iHPCs, and primary cord-blood (CB) derived CD34+ cells (GSE40799) (Weidner et al., 2013). **(c)** Epigenetic age was predicted with Horvath's clock (Horvath et al., 2018) for iPSCs, iMSCs, and iHPCs. **(d)** Canyons were analyzed in the genes *PAX6*, and *GATA2* for iPSCs, iMSCs, and iHPCs. The two exemplary genes were selected for comparison to Jeong et al., 2018.

### Supplemental tables

**Table S1: gRNAs for CRISPR.**

| Primer | Sequence (5' – 3') |
|---|---|
| Exon 2_1a FW | CACCGACAGAGGCACCGTTCACCAG |
| Exon 2_1a RV | AAACCTGGTGAACGGTGCCTCTGTC |
| Exon 2_1b FW | CACCGCCTGTGGGTGGGGGCTTCGA |
| Exon 2_1b RV | AAACTCGAAGCCCCCACCCACAGGC |
| Exon 2_2a FW | CACCGATCACTCAGTGCTCACGACA |
| Exon 2_2a RV | AAACTGTCGTGAGCACTGAGTGATC |
| Exon 2_2b FW | CACCGGCGGTCATGCACTCAGTATG |
| Exon 2_2b RV | AAACCATACTGAGTGCATGACCGCC |
| Exon 19_1a FW | CACCGTTTCTCTTCCGACCTCTCAG |
| Exon 19_1a RV | AAACCTGAGAGGTCGGAAGAGAAAC |
| Exon 19_1b FW | CACCGCAGCTGGGGCTGTCTGCAT |
| Exon 19_1b RV | AAACATGCAGACAGCCCCAGCTGC |
| Exon 19_2a FW | CACCGGGACATCTCGCGATTTCTCG |
| Exon 19_2a RV | AAACCGAGAAATCGCGAGATGTCCC |
| Exon 19_2b FW | CACCGCCTCTTGTCACTAACGCCCA |
| Exon 19_2b RV | AAACTGGGCGTTAGTGACAAGAGGC |
| Exon 23_1a FW | CACCGTAGACGGCTTCCGGGCAGCC |
| Exon 23_1a RV | AAACGGCTGCCCGGAAGCCGTCTAC |
| Exon 23_1b FW | CACCGAACCACACAGCAGGACCCGG |
| Exon 23_1b RV | AAACCCGGGTCCTGCTGTGTGGTTC |
| Exon 23_2a FW | CACCGCTTTGCCTTGCGAAAAGGGT |
| Exon 23_2a RV | AAACACCCTTTTCGCAAGGCAAAGC |
| Exon 23_2b FW | CACCGCTCCGGTATTTCCGCCTCTG |
| Exon 23_2b RV | AAACCAGAGGCGGAAATACCGGAGC |

**Table S2: Flanking primers for PCR.**

| Primer | Sequence (5' – 3') |
|---|---|
| Exon 2 FW | GGTTTCTCTGCTCTCTGGGGTG |
| Exon 2 RV | CTGGAACCAAATGCCTGCTCC |
| Exon 19 FW | GCAATGACCTCTCCATCGTCA |
| Exon 19 RV | CCCTGCTTCCTCCCTTTCTATC |
| Exon 23 FW | GAAACCAGCACAGTGCCTGG |
| Exon 23 RV | GAAGGGGGAGGAAGGGAA |

**Table S3: Exon specific primers for qPCR.**

| Primer | Sequence (5' – 3') |
|---|---|
| Exon 2-3 FW | CGGGGACACCAGCAGCTC |
| Exon 2-3 RV | GCTTGCGCTTCCTCCCAG |
| Exon 16-17 FW | GGTGCTGTCTCTCTTTGATGGA |
| Exon 16-17 RV | CCGACGTACATGATCTTCCCC |
| Exon 17/18-19 FW | GAAGCATATCCAGGAGTGGGG |
| Exon 17/18-19 RV | CAGGAGGCGGTAGAACTCAAA |
| Exon 22-23 FW | AGGGCAAAGACCAGCATTTTC |
| Exon 22-23 RV | CCAAGCGGCTCATGTTGGAG |
| GAPDH FW | TTCGTCATGGGTGTGAACCA |
| GAPDH RV | CTGTGGTCATGAGTCCTTCCA |

**Table S4: qPCR primers for the trilineage assay.**

| Primer | Sequence (5' – 3') |
|---|---|
| OCT3/4 FW | GGGGGTTCTATTTGGGAAGGTA |
| OCT3/4 RV | ACCCACTTCTGCAGCAAGGG |
| NANOG FW | CAGAAGGCCTCAGCACCTAC |
| NANOG RV | ATTGTTCCAGGTCTGGTTGC |
| CD34 FW | TGGACCGCGCTTTGCT |
| CD34 RV | CCCTGGGTAGGTAACTCTGGG |
| NKX2.5 FW | ACCTCAACAGCTCCCTGACTCT |
| NKX2.5 RV | ATAATCGCCGCCACAAACTCTCC |
| MYH6 FW | AAGCTCAAGAACGCCTAC |
| MYH6 RV | CATTCTTTCCTCCTTCTCC |
| AFP FW | GCCAAGCTCAGGGTGTAG |
| AFP RV | CAATGACAGCCTCAAGTTGT |
| ALB FW | GGTGTGTTTCGTCGAGATG |
| ALB RV | ACTGAGCAAAGGCAATCAAC |
| CD31 FW | GAGTCCTGCTGACCCTTCTG |
| CD31 RV | ATTTTGCACCGTCCAGTCC |
| FOXA2 FW | GCATTCCCAATCTTGACACGGTGA |
| FOXA2 RV | GCCCTTGCAGGCAGAATACACATT |
| SOX17 FW | AGGAAATCCTCAGACTCCTGGGTT |
| SOX17 RV | CCCAAACTGTTCAAGTGGCAGACA |
| GFAP FW | AGGAGGAGGTTCGGGAACTC |
| GFAP RV | CGCCATTGCCTCATACTGC |
| NESTIN FW | CCTCAAGATCTCCCTCAGCC |
| NESTIN RV | CCAGCTTGGGGTCCTGAAAG |
| PAX6 FW | TCGAAGGGCCAAATGGAGAAGAGAAG |
| PAX6 RV | GGTGGGTTGTGGAATTGGTTGGTAGA |
| SOX1 FW | CCTGTGTGTACCCTGGAGTTTCTGT |
| SOX1 RV | TGCACGAAGCACCTGCAATAAGATG |
| GAPDH FW | GAAGGTGAAGGTCGGAGTC |
| GAPDH RV | GAAGATGGTGATGGGATTTC |

**Table S5: Differentially methylated CpGs in iPSC WT *versus DNMT3A* knockouts.**

This table is provided as Supplemental file 2.xlsx

**Table S6: Differentially methylated CpGs in iPSCs *versus* iMSCs (WT or *DNMT3A* knockouts).**

This table is provided as Supplemental file 3.xlsx

**Table S7: Differentially methylated CpGs in iPSCs *versus* iHPCs (WT or *DNMT3A* knockouts).**

This table is provided as Supplemental file 4.xlsx

**Table S8: Differentially methylated CpGs in iHPC WT *versus DNMT3A* knockouts.**

This table is provided as Supplemental file 5.xlsx

**Table S9: RNA-seq data of iHPCs.**

This table is provided as Supplemental file 6.xlsx

### Supplemental experimental procedures

#### Western blot

To validate *DNMT3A* knockouts, cells were lysed with RIPA buffer and 60 µg protein lysate was loaded onto 12% Mini-PROTEAN TGX Precast Protein Gels (BioRad, Hercules, California, USA). Membranes were blocked with 4% skim milk (Sigma-Aldrich) in TBST for 1 h at room temperature and then incubated with the primary DNMT3A antibody (#2160, Cell Signaling Technology, Danvers, MA, USA) and with the β-actin antibody (clone AC-74, Sigma Aldrich, St. Louis, MO, USA) at 4°C overnight. As secondary antibodies Peroxidase AffiniPure Goat Anti-Rabbit IgG (H+L) (Jackson Immuno Research, West Grove, Pennsylvania, USA) and Goat IgG anti-mouse IgG (H+L)-HRPO (Dianova, Hamburg, Germany) were used and incubated for 1 h at room temperature. Detection was performed with the Super Signal West Dura Extended Duration Substrate (Thermo Fisher Scientific) and X-ray films (CL-XPosure, Thermo Fisher Scientific) with the Optimax 2010 x-ray film processor (Protec, Oberstenfeld, Germany).

#### Real-time PCR

For semiquantitative real-time PCR (qRT-PCR), RNA was isolated with the NucleoSpin RNA Plus kit (Macherey Nagel). 500 ng RNA was reverse transcribed using the High-Capacity cDNA Reverse Transcription Kit (Thermo Fisher Scientific) and amplified with the StepOnePlus Real-Time PCR system using the Power SYBR Green PCR Master Mix (both Thermo Fisher Scientific) with specific primers (Tab. S3,S4). Expression levels were normalized to *GAPDH*.

#### Immunofluorescence

For immunofluorescence staining, iPSCs were seeded onto glass slides coated with vitronectin, cultivated until reaching a sufficient colony size, fixed with 4% paraformaldehyde, blocked with normal goat serum for 30 minutes and stained with TRA-1-60 antibody (clone TRA-1-60, Merck Millipore, Burlington, MA, USA) or OCT4 antibody (clone sc-9081, Santa Cruz Biotechnology, Dallas, TX, USA) over night at 4°C. As secondary antibodies goat anti-mouse IgM Alexa Fluor 594 and goat anti-rabbit IgG FITC (both Thermo Fisher Scientific) were used respectively. Nuclei were counterstained with DAPI for 15 minutes. Microscopic pictures were taken with the fluorescence microscope Axioplan 2 (Carl Zeiss, Oberkochen, Germany).

#### Differentiation in embryoid bodies

For spontaneous trilineage differentiation, iPSCs were seeded in three different densities (1 500; 3 000 and 5 000 cells per well) into U-bottom 96-well plates (TPP, Trasadingen, Switzerland). Plates were centrifuged for 5 minutes at 350 x g and then incubated at 37° C and 5% $CO_2$ in iPS Brew XF with ROCK-inhibitor and 0.4% polyvinylalcohol (PVA). After 24 h, medium was changed to EB-medium consisting of 77% KO-DMEM, 5% KO serum replacement, 100 U/mL penicillin, 100 µg/mL streptomycin, 1% non-essential amino acids and 0.5% β-mercaptoethanol (all Thermo Fisher Scientific) with daily medium changes. After 7 days EBs were transferred to 0.1% gelatin coated 6-well plates with in total around 30 EBs per well and cultured until day 28.

#### Immunophenotypic analysis

Flow cytometric analysis was performed on a FACS Canto II (BD Biosciences, Franklin Lakes, New Jersey, USA) and analyzed with FlowJo software Version 10.4.2 (FlowJo LLC, Ashland, Oregon, USA). The following antibodies were used for iMSC analysis: CD14-APC (clone M5E2), CD29-PE (clone MAR4), CD31-PE (clone WM59), CD34-APC (clone 581), CD45-APC (clone HI30), CD73-PE (clone AD2), CD90-APC (clone 5E10) (all BD Biosciences), and CD105-FITC (clone MEM-226; ImmunoTools, Friesoythe, Germany). For analysis of iHPCs we used CD3-APC (clone HIT3a), CD31-PE (clone WM59), CD34-APC (clone 581), CD43-FITC (clone 1G10), CD66b-PE (clone G10F5) (all BD Biosciences, Franklin Lakes, USA), CD11c-PE-Cy7 (clone 3.9), HLA-DR-FITC (clone LN3), CD235a-PE (clone HIR2/GA-R2), cKIT-PE-Cy7 (clone 104D2), (all Thermo Fisher Scientific), CD45-APC-Vio770 (clone 5B1), CD33-APC (clone AC104.3E3) (all Miltenyi Biotec), and CD61-FITC (clone VI-PL2, Biolegend, San Diego, CA, USA).

#### Colony forming unit assay

After 16 days of differentiation 5 000 iHPCs were seeded in 500 µL of methylcellulose based medium (HSC-CFU lite with EPO; Miltenyi Biotec) in 24-well plates. Colonies were scored after two weeks of culture.

#### Competitive hematopoietic differentiation using RGB LeGO vectors

Wildtype, exon $19^{-/-}$ and exon $23^{-/-}$ iPSCs were transduced with Venus, Cerulean, or mCherry RGB libraries, respectively. To this end, iPSCs were grown until a confluency of 30-40%. 200 µL of virus suspension supplemented with 8 µg/mL polybrene (Sigma-Aldrich, St. Louis, MO, USA) was added to the culture medium and cells were centrifuged in 6-well plates at 480 rcf for 2 h and incubated for another hour at 37°C and 5% $CO_2$. Subsequently, virus suspension was replaced with fresh iPS-Brew XF (Miltenyi Biotec). After 24 h, transduction was repeated by incubation of cells with the virus/polybrene mix for 3 h at 37°C and 5% $CO_2$. Puromycin selection was started 24 h post-transduction for 2-3 days with increasing puromycin concentrations from 1.0 µg/mL to 1.5 µg/mL. After a short expansion phase the barcoded wildtype, exon $19^{-/-}$ and exon $23^{-/-}$ iPSCs (all donor 1 with same genetic background) were mixed at equal cell numbers for a competitive hematopoietic differentiation assay for 15 days. Thereafter, iHPCs were expanded for additional 28 days in StemSpan SFEM (Stemcell Technologies) supplemented with 10 ng/mL fibroblast growth factor 1 (FGF-1), 10 ng/mL SCF, 10 ng/mL TPO (all purchased from Peprotech), 100 U/mL penicillin and 100 µg/mL streptomycin. Flow cytometric analysis of RGB cells was performed with a Cytek Aurora (Cytek Biosciences, Fremon, CA, USA) and analyzed with the FlowJo software Version 10.4.2. Fluorescence images of RGB cells were acquired with a Zeiss Axiovert 200 (Carl Zeiss, Oberkochen, Germany).
